# Zebrafish models of candidate human epilepsy-associated genes provide evidence of hyperexcitability

**DOI:** 10.1101/2024.02.07.579190

**Authors:** Christopher Mark LaCoursiere, Jeremy F.P. Ullmann, Hyun Yong Koh, Laura Turner, Cristina M. Baker, Barbara Robens, Wanqing Shao, Alexander Rotenberg, Christopher M. McGraw, Annapurna Poduri

**Affiliations:** F.M. Kirby Neurobiology Center, Department of Neurology, Boston Children’s Hospital, Boston, MA, USA; Epilepsy Genetics Program, Department of Neurology, Boston Children’s Hospital, Boston, MA, USA; Departments of Neuroscience and Pediatrics, Division of Neurology and Developmental Neuroscience, BCM, Houston, Texas, USA; Kavli Institute for Systems Neuroscience, Norwegian University of Science and Technology, Trondheim, Norway; Research Computing, Department of Information Technology, Boston Children’s Hospital, Boston, MA, USA; Department of Neurology, Harvard Medical School, Boston, MA, USA; Division of Epilepsy and Clinical Neurophysiology, Department of Neurology, Boston Children’s Hospital, Boston, MA, USA; Department of Neurology, Massachusetts General Hospital, Boston, MA, USA; Broad Institute of Harvard and MIT, Cambridge, MA, USA

## Abstract

Hundreds of novel candidate human epilepsy-associated genes have been identified thanks to advancements in next-generation sequencing and large genome-wide association studies, but establishing genetic etiology requires functional validation. We generated a list of >2200 candidate epilepsy-associated genes, of which 81 were determined suitable for the generation of loss-of-function zebrafish models via CRISPR/Cas9 gene editing. Of those 81 crispants, 48 were successfully established as stable mutant lines and assessed for seizure-like swim patterns in a primary F_2_ screen. Evidence of seizure-like behavior was present in 5 (*arfgef1, kcnd2, kcnv1, ubr5, wnt8b*) of the 48 mutant lines assessed. Further characterization of those 5 lines provided evidence for epileptiform activity via electrophysiology in *kcnd2* and *wnt8b* mutants. Additionally, *arfgef1* and *wnt8b* mutants showed a decrease in the number of inhibitory interneurons in the optic tectum of larval animals. Furthermore, RNAseq revealed convergent transcriptional abnormalities between mutant lines, consistent with their developmental defects and hyperexcitable phenotypes. These zebrafish models provide strongest experimental evidence supporting the role of *ARFGEF1, KCND2*, and *WNT8B* in human epilepsy and further demonstrate the utility of this model system for evaluating candidate human epilepsy genes.

**Highlights:**

- Zebrafish models generated by CRISPR/Cas9 gene editing display seizure-like swim patterns in five candidate human epilepsy genes: *arfgef1, kcnd2, kcnv1, ubr5, wnt8b*.
- Local field potential abnormalities recorded from *kcnd2* and *wnt8b* crispants provide additional evidence of hyperexcitability.
- *Arfgef1* and *wnt8b* mutant larvae have fewer inhibitory interneurons than wild type in the optic tectum.
- CRISPR-generated mutants of epilepsy genes displayed convergent transcriptional dysregulation, consistent with developmental abnormalities and their hyperexcitability phenotype.

## Introduction

Epilepsy is a condition defined by recurrent, unprovoked seizures affecting 1% of the population, including 1 in 200 children[1], with a lifetime prevalence of 1 in 26[2]. Major personal and societal impacts for people living with epilepsy include refractory seizures, neurodevelopmental and neuropsychiatric comorbidities, high epilepsy-related healthcare costs, and increased risk of sudden death[3]. Underlying etiologies are heterogeneous and include both acquired and genetic factors. Recent technological advancements in genetics research have enabled the characterization of numerous monogenic epilepsies[4-6], collectively accounting for a substantial portion of otherwise clinically unexplained disorders[7]. Through collaborative efforts to study the role of genetics in the epilepsies, as well as those of individual laboratories, the number of genes associated with epilepsy has steeply risen to >2,000[5, 6, 8-10]. While a few well-powered, large studies have securely implicated novel genes in early onset epilepsy, many reports have revealed candidate genes that require validation[11], both in additional patient cohorts and in experimental systems. Despite major progress in gene discovery, treatment for epilepsy continues to be empiric and often unsuccessful, with at least one-third of patients with epilepsy continuing to experience seizures despite treatment with anti-seizure medications (ASMs). The development of rational treatments for monogenic epilepsies requires clinically relevant functional models to dissect gene-specific epileptogenic mechanisms and develop targeted therapies[12].

Characterization of the numerous, heterogeneous genetic epilepsies poses a challenge, as many distinct models are needed to study the wide range of biological processes that, when disrupted, result in circuit dysfunction and hyperexcitability. An additional challenge stems from the fact that these models of central nervous system (CNS) dysfunction may result in broad neurodevelopmental phenotypes or potentially only transient episodic phenotypes akin to human seizures[13]. Overcoming these challenges requires *in vivo* models and multiple diverse modalities to characterize normal and disease states. Conventional epilepsy models, such as rodent, are expensive and time-consuming to generate and, further, are not guaranteed to display phenotypes relevant to human epilepsy, including the most basic phenotype of seizures[14]. Zebrafish (*Danio rerio*) are an increasingly employed epilepsy model because of their high genetic homology to humans, large clutch sizes, inexpensive husbandry costs, and the ease with which their small, transparent larvae readily allow for morphologic phenotyping[15]. The relevance of zebrafish to epilepsy disease modeling and drug discovery has been demonstrated by the presence of seizure-like events and electrophysiological abnormalities similar to those observed in human epilepsy[16], as well as translation from drug screens in genetic models of *SCN1A*-related epilepsy, a prototypical genetic epilepsy with childhood onset and associated cognitive comorbidities and risk of sudden death[17].

Motivated by burgeoning human genetics discoveries, we present the results of a literature-based search for novel genes associated with developmental epilepsies, assessment for suitability of modeling in zebrafish, CRISPR model generation for 81 genes, and automated seizure detection followed by validation and characterization of the 5 most promising models. Our pipeline and results further the priorities of the epilepsy genetics community for pre-clinical, precision medicine approaches to diagnose and treat genetic epilepsies.

## Results

### Selection of candidate human epilepsy-associated genes for modeling in zebrafish

To select genes suitable for modeling epilepsy in zebrafish, we first generated a master list of >2,200 unique epilepsy-related genes consolidated and vetted by the neurologists and genetic counselors of our Epilepsy Genetics Program (Figure 1A.i). This list was refined to include genes that are intolerant to missense variation (Z-score>3) or loss-of-function (LOF) (probability of LOF intolerance, pLI>0.9) according to metrics from the Exome Aggregation Consortium database[18] and subsequently the gnomAD database[19]. Final *in silico* refinement included genes with zebrafish-human homology of at least 60% with no previously established animal model. A small selection of candidates not meeting all metrics, but with additional evidence of pathogenicity from exome data analyzed through the Epilepsy Genetics Program, were also included (Figure 1Aii, Supplemental Table 1). This resulted in a list of 81 genes with highly heterogenous molecular functions (Figure 1B, Supplemental Table 1) predicted to produce LOF models with protein-coding insertions and deletions (indels) confirmed via Synthego ICE analysis (https://ice.synthego.com). Given preliminary >90% cutting efficiency, a shotgun gene targeting approach[20], and a gene list highly intolerant to LOF, any transcript that failed to transmit (>8 pools of 5 F_1_ larvae) to F_1_ germline was assumed likely embryonic lethal. Additionally, F_1_ fish that did not result in viable heterozygous (HET) in-cross (after >10 breeding opportunities) and thus did not provide F_2_ offspring for downstream analysis were not pursued further. In total, 48 candidates led to viable models subjected to further evaluation (Figure 1A.iii). Functional validation of suitable candidates began with three larval swim paradigms (detailed in methods), which implicated 5 of 48 mutants with classic larval zebrafish seizure (stage SII) swim patterns (Figure 1A.iv). Indel mutations in those 5 lines (*arfgef1, kcnd2, kcnv1, ubr5*, and *wnt8b)* are depicted in Figure 1C. Notably, 3/5 mutations were in-frame and not predicted to result in truncation or complete KO as confirmed by Sanger sequencing. *Arfgef1* models contain a 10 base pair deletion and 4 base pair insertion resulting in a p.L367S missense as well as a two amino acid deletion at p.E368_E369. *Kcnd2* editing resulted in an in-frame 3-base pair deletion at the evolutionarily conserved amino acid p.M375 (Figure 1C), while *wnt8b* mutants contain a ten base pair deletion resulting in splice site removal and deletion of two amino acids at p.S79_R80. Indels in *kcnv1* and *ubr5* models are early truncating frameshift mutations. (Figure 1C, Supplemental Table 2). These five behaviorally positive mutants were then subjected to downstream assays to provide additional evidence of hyperexcitability. (Figure 1A.v).

**Figure 1.**
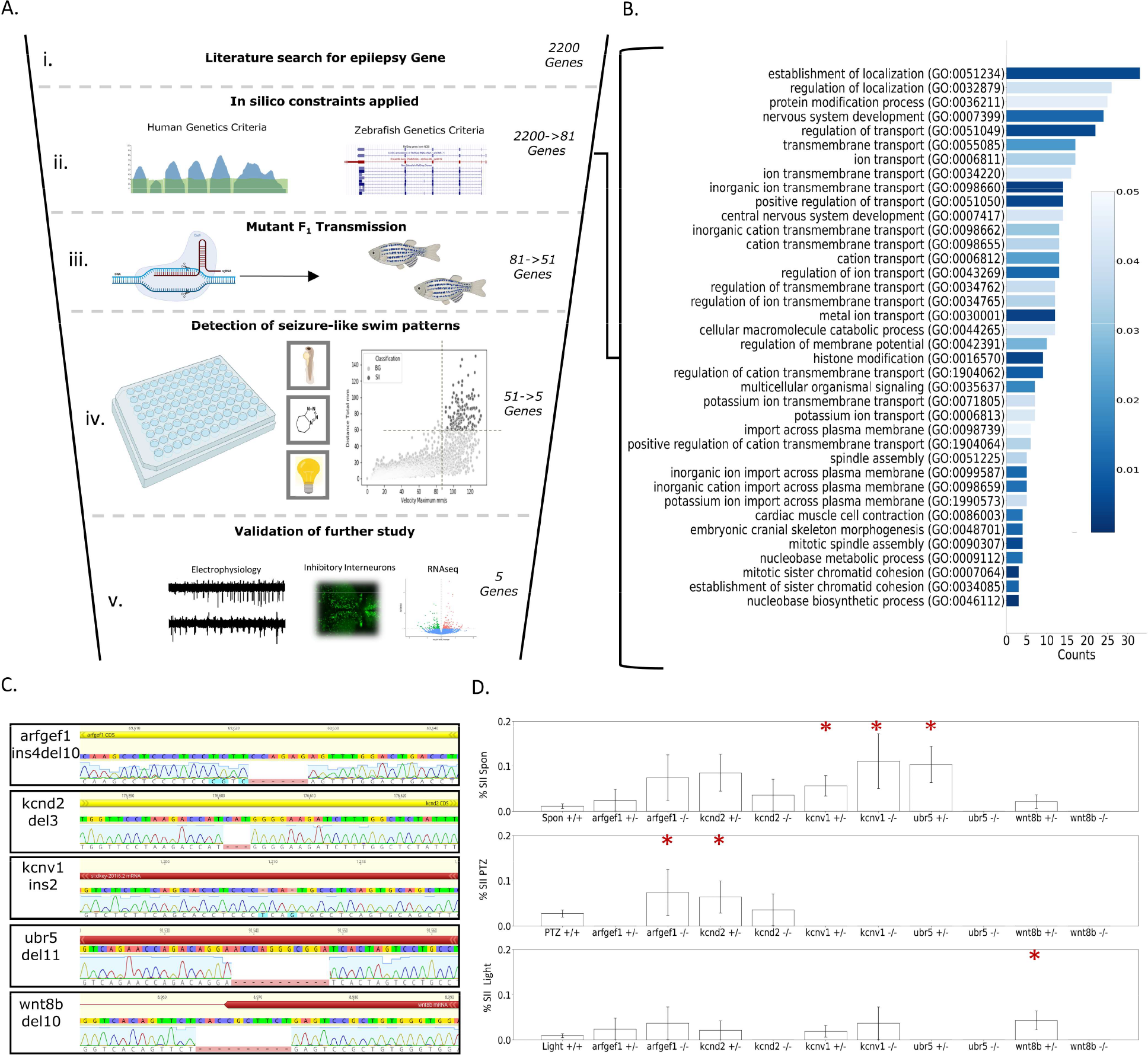
Overview of evaluation of candidate human epilepsy genes in zebrafish models. (A) Graphical depiction of primary F_2_ screening method including: i. initial gene identification through literature search; ii. application of in silico constraints; iii. generation of initial crispants and stable F2 mutants; iv. detection of seizure-like swim patterns in either a spontaneous, light-provoked, or subthreshold PTZ assay, v. and characterization of mutant lines that displayed SII seizure events via local field potentials, quantifying *dlx*-GFP expressing inhibitory interneurons and RNAseq. (B) 48 genes for which we generated surviving crispants and stable mutant lines represent heterogeneous Gene Ontology (GO) categories of biological processes. (C) Schematic of homozygous mutations found in the five genes meeting criteria for positive seizure-like swim patterns. (D) Results from hyperexcitable behavior positive larvae assayed in three paradigms. Paradigms included a 1hr spontaneous recording, a light provoked recording and a subthreshold PTZ provoked paradigm. Mutant larvae exhibited limited SI-like behavior as shown by mean velocity, but 5 mutant alleles showed an increase in the proportion of larvae with at least one SII event. Statistical significance for SI behavior was determined by Kruskal-Wallis tests with Dunn multiple comparisons while significance of SII events was determined by the binomial probability of SII events compared to the expected value from the population proportion. Significance was determined within a 95% confidence interval, \**p <*0.05, \*\**p <*0.01, \*\*\**p <*0.001. Figure (A) was created with BioRender.com.

### Seizure-like swim patterns provide initial evidence of gene-disease association in 5 mutant lines

Initial F_2_ screening was conducted by recording and analysis of locomotion with analysis of swim patterns for seizure-like patterns at 5dpf. No obvious increase in normal beat and glide swim (SI) activity was observed in any ungenotyped mutant assays (Supplemental Table 2). Previous studies[15] have shown that classic swirling swim patterns representing seizure events (SII) can be followed by a postictal-like decrease in movement. Detection of SII events may be limited using average velocity to screen for abnormal behavior due to periods of prolonged inactivity following the seizure events. Therefore, we developed parameters to identify discrete SII events, and SII detection was automated from video files with a classifier algorithm (detailed in methods). A “positive” SII result for a given gene indicates an increased binomial probability of SII events in mutant populations compared to the wildtype (WT) SII population. We observed that 5 candidate gene models displayed statistically increased probability of SII events. Of the five genes with increased SII activity *wnt8b* was the only mutant that showed an abnormality in SI swim pattern, which was in fact a decrease (Supplemental table 3).

We observed evidence of seizure events in 5 mutant lines: *arfgef1, kcnd2, kcnv1, ubr5*, and *wnt8b*. Specifically, we observed increased binomial probability of SII seizure events in mutants vs. WT larvae in at least one of three paradigms (spontaneous, light-induced, subthreshold PTZ-induced). Both *kcnv1* HET and *kcnv1* homozygous (HOM) larvae displayed spontaneous seizures, with a total of 19 SII seizure events across all paradigms (Figure 1D). There was a total of 8 SII seizure events in *kcnd2* HET larvae across all paradigms. *Arfgef1* HOM mutants showed a significant increase in light-induced SII seizure events (average 8 total SII events vs. 0 in WT). *Ubr5* HET mutants, but interestingly not *ubr5* HOM mutants, displayed increased spontaneous seizure-like events vs. WT. Similarly, *wnt8b* HOM mutants displayed light paradigm induced SII seizure (Figure 1D). We noted that several WT larvae generated from both *wnt8b* and *ubr5* crosses had higher tendency to show SII seizures than stock WT data (Supplemental Table 2), suggesting the need for comparisons to be made with conspecific WT larvae. Together these data demonstrate that SII behavioral seizures occur only rarely as assessed by our experimental paradigms, and they implicate 5 genes from our primary F_2_ screen for further characterization.

### Additional characterization of hyperexcitability

Of the 5 genes demonstrating seizures using high-throughput detection of SII swim patterns, 2 had models that additionally displayed abnormal and hyperexcitable electrophysiological activity. Recordings were conducted in the optic tectum, a conventional location for recording that allows demonstration of epileptiform activity in larval zebrafish[21-23](Figure 2A,B,C). The candidate genes were assayed (HET and HOM mutants) and compared to conspecific WTs 5-days post fertilization (dpf) during a 22-minute window of a half-hour recording to allow for initial acclimatization. Mutant larvae displayed a range of abnormal spiking including an increase in spike count and spike amplitude (Figure 2D,E). We observed a wide range of epileptiform-like values in WT groups in all parameters assayed, consistent with the necessity for conspecific comparisons in LFP experiments as previously described[21]. *Kcnd2* mutants proved to have the strongest evidence of hyperexcitability with an increase in average spike rate and amplitude as well as increased bursting duration and frequency. All parameters indicated a dose dependence, with the severity of phenotypes increasing from HET to HOM. *Wnt8b* mutants also exhibited a dose-dependent increase in hyperexcitability in spike rate, burst count, and burst duration, with homozygous mutants more severely affected than heterozygous clutch mates (Figure 2F,G). The *arfgef1* (Figure 2G-E), *ubr5*, and *kcnv1* mutants did not display statistically significant abnormalities in LFP measurements (Supplemental Table 3).

**Figure 2.**
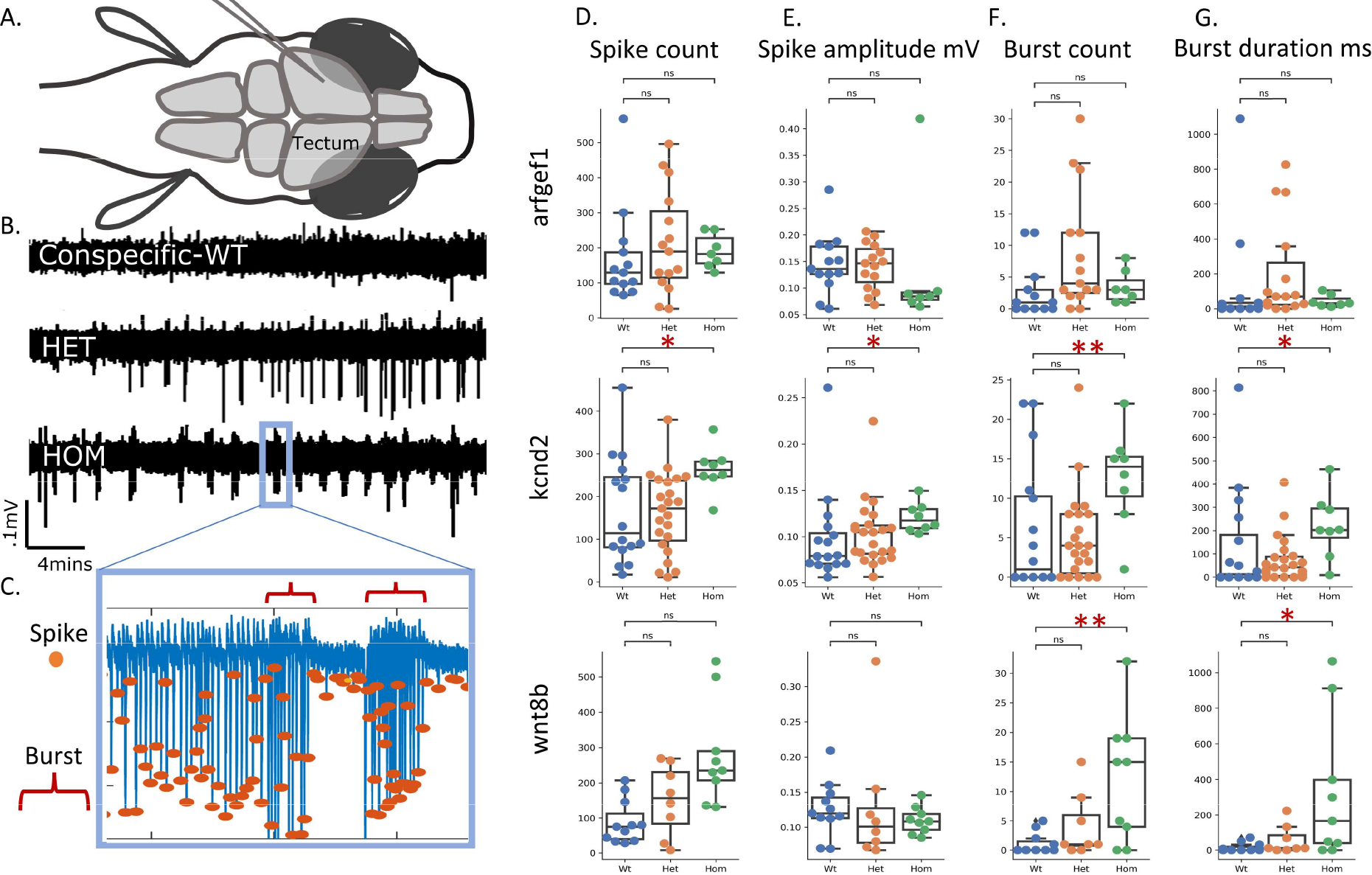
Epileptiform activity of larval tectum in electrophysiological recordings. (A) Schematic representation of larval zebrafish LFP recordings. (B) Exemplary LFP recordings from kcnd2 larvae showing a dose sensitive increase in spike amplitude, frequency, distribution, and bursting. (C) Exemplary recordings of spike count, amplitude and burst count and duration. (D) Spike count quantification is increased *kcnd2* and *wnt8b* HOM larvae compared to conspecific controls. (E) Spike amplitude is increased in HOM kcnd2 larvae compared to conspecific control. (F and G) Burst count and duration increased in *kcnd2* and *wnt8b* HOM compared to conspecific controls. Statistical significance was determined by Kruskal-Wallis test with Dunn multiple comparisons, \**p <* 0.05, \*\**p <* 0.01, \*\*\**p <* 0.001.

### Reduced interneuron number in candidate epilepsy gene models

To assess whether abnormal larval tectal development could contribute to the SII seizures and hyperexcitable phenotypes observed in LFP experiments, we quantified inhibitory interneurons in WT, HET, and HOM larvae (Figure 3A). No qualitative difference in the distribution of inhibitory interneurons was observed between mutant and WT forebrain, tectum, and hindbrain of age-matched 5 dpf larvae. Dense clusters aggregated near anatomical boundaries and the midline and dispersed radially with no gross differences between mutants and conspecific controls (Figure 3B). Regional quantification of larval tectum was isolated and quantified to directly probe development of observed hyperexcitable structures. Of the 3 candidate genes with increased excitation from LFP experiments, 2 showed a significant reduction in inhibitory interneurons: *arfgef1* and *wnt8b*. Both genes’ models showed a decrease in HET interneuron quantity in the larval tectum followed by a more dramatic decrease in HOM larvae (Figure 3C). *Kcnv1* and *kcnd2* models showed no indication of GABAergic interneuron reduction (Supplemental Table 3).

**Figure 3.**
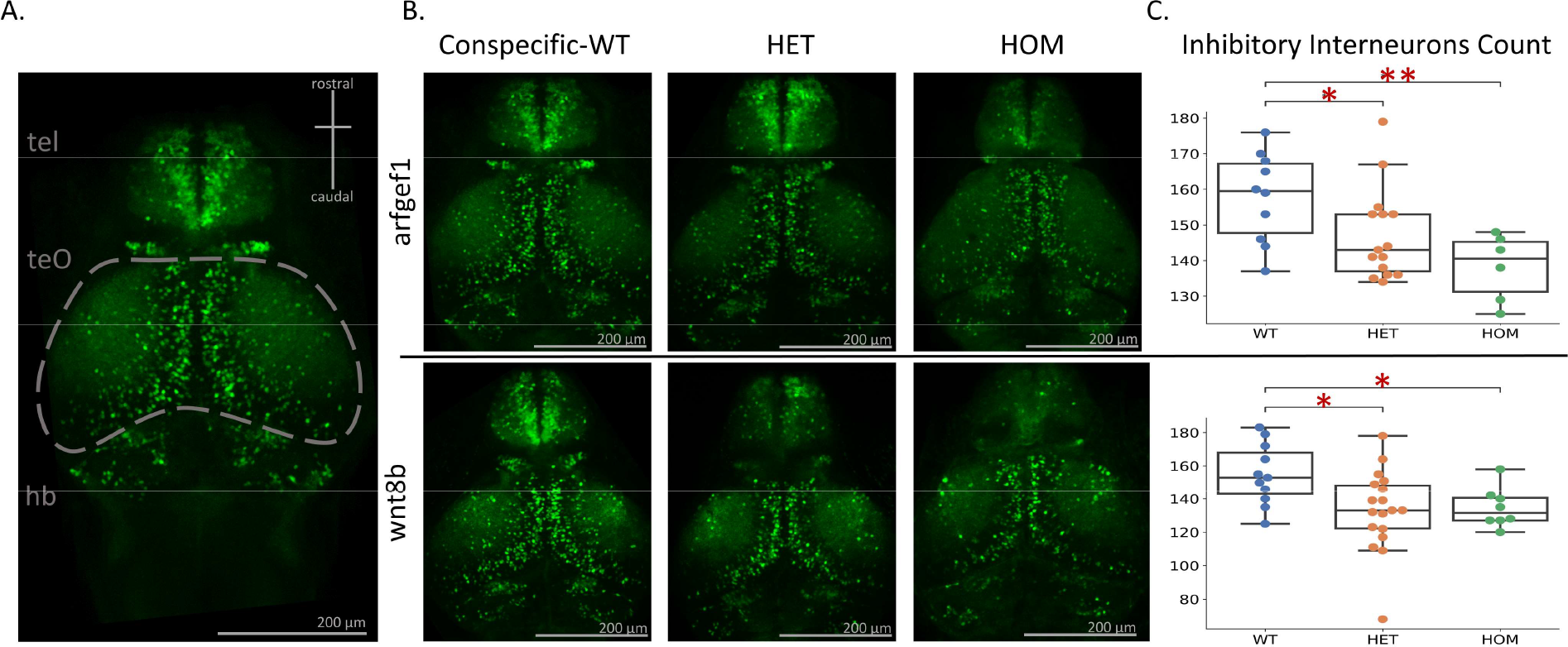
A dose-dependent decrease in the number of interneurons in a subset of zebrafish mutant lines, *arfgerf1* and *wnt8b*. (A) Schematic representation of larval zebrafish brain gross anatomy highlighting the optic tectum. (B) Exemplary images of *arfgef1* and *wnt8b* mutant allele combinations. (C) Dose dependent decrease in inhibitory interneurons in *arfgef1* and *wnt8b* mutants compared to conspecific controls. tel, telencephalon; teO, optic tectum; hb, hindbrain. Statistical significance was determined by Kruskal-Wallis test with Dunnett’s multiple comparisons, \**p <* 0.05, \*\**p <* 0.01, \*\*\**p <* 0.001.

### Transcriptomic differences in zebrafish epilepsy models

Given the heterogeneity of genetic causes of epilepsy but the convergence of a hyperexcitability phenotype, transcriptomic patterns were assessed across the mutant zebrafish that showed structural or electrophysiological abnormalities to probe for convergent pathways that might lead to hyperexcitability. Larval heads were dissected and pooled by genotype and assayed via RNA-Seq (Supplemental Figure 1A). We identified thousands of differentially expressed genes (DEGs) within mutants as compared to conspecific controls (Figure 4A.i-A.iii). The most striking differences were seen in candidates with strong findings via LFP; *kcnd2* and *wnt8b* (Figure 4A.ii-Aiii). Directionality of dysregulated genes within HET and HOM conspecifics for the same gene were largely overlapping and consistent (Figure 4A.i-A.iii). Considering DEGs for only Hom vs WT for each mutant line, data from *wnt8b* mutant larvae displayed the most dysregulation, followed by *kcnd2* and finally *arfgef1* (Figure 4B.i-B.iii). Only the mutant *arfgef1* HOMs show a significant reduction in mutant gene transcription as compared with WT (Figure 4A.i). Of those genes that were dysregulated in mutant lines; DEG co-occurrence from our previously published EPGP epilepsy gene list[10] was seen in the top twenty most significantly dysregulated genes in 2 HOM mut lines. These genes include, *atp8a2* in *arfgef1* mutants (Figure 4C.i) as well as *col4a1* and *grin1b* in *wnt8b* mutants (Figure 4C. iii). When considering all co-occurrence between mutant line DEGs and the EPGP gene list, the highest number was seen in *kcnd2* (n = 5/1443) and *wnt8b* (n = 298/1443) mutants. However, when the total number of DEGs were taken into consideration, we did not observe an enrichment of the EPGP gene list within the DEGs, nor did we observe an enrichment of DEGs within the EPGP gene list for any mutants (Supplemental Figure 1B). Interestingly, we observed transcriptional biomarkers for neuronal activity, immediate early genes (IEG), in the mutant lines with hyperexcitability LFP recordings. Dysregulation of *npas4a, npas4b, erg1, fosab*, and *nr4a1* was seen in mutant *wnt8b* (Figure 4D.ii) fish and *fosab, erg1*, and *npas4b* in *kcnd2* (Figure 4D.ii) mutants. All IEG expression was significantly downregulated, consistent with chronic epilepsy as opposed to the upregulation observed after acute seizure induction[24-26]. Models with no significant findings from LFP also showed no significant IEG DEGs (Figure 4C, D,E.ii). The highest number of co-occurrences of DEGs between mutant lines was also seen between *kcnd2* and *wnt8b* mutants while minimal co-occurrence was seen between *arfgef1* and *kcnd2* or *wnt8b* (Supplemental Figure 1C).

**Figure 4.**
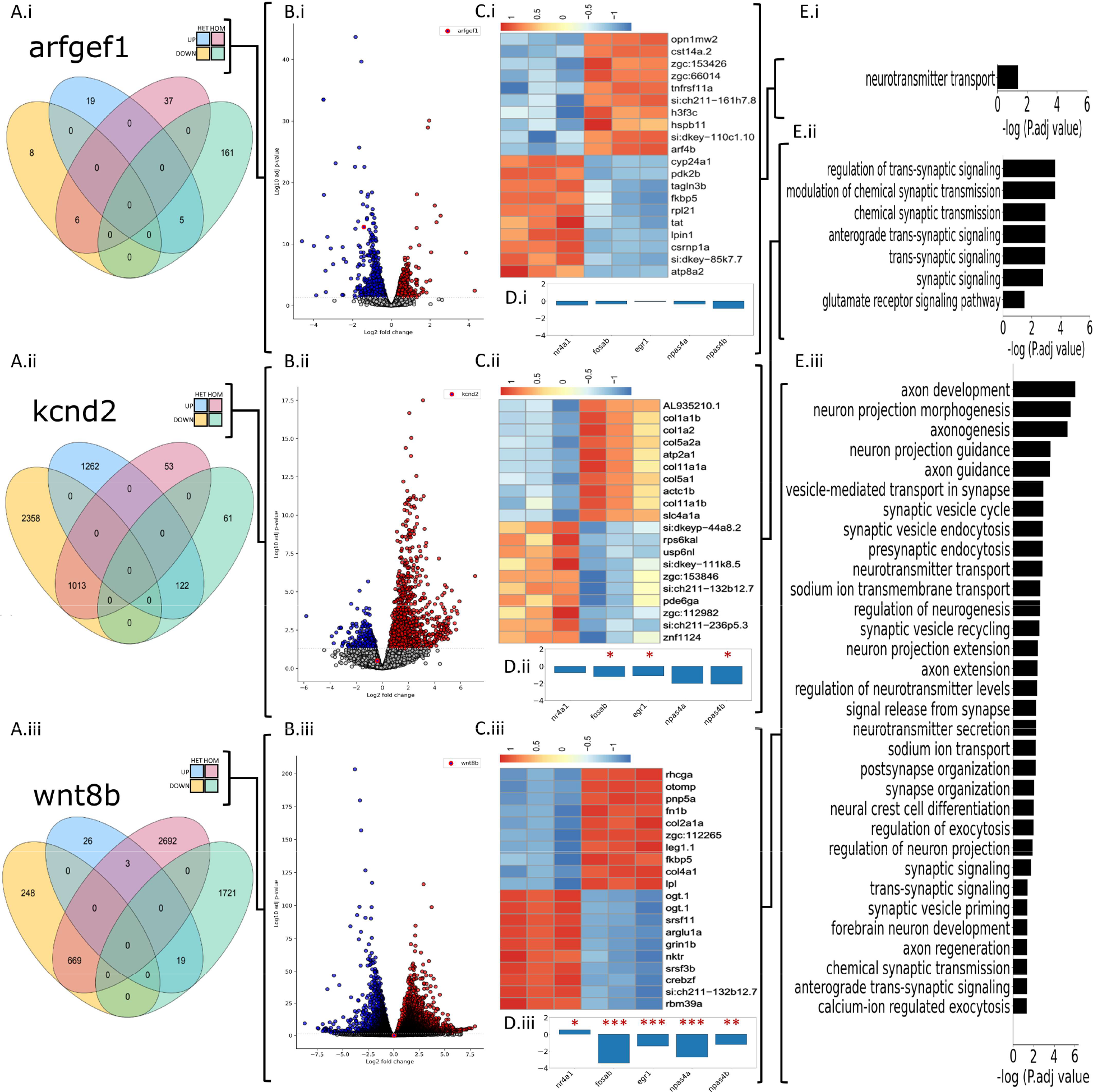
Mutants show dysregulation and coincidence, among phenotypically similar mutants, of epilepsy associated genes and ontologies. (A.i-A.iii) Venn diagram of the total number differentially expressed genes (DEGs) per mutant genotype. (B.i-B.iii) Volcano plot of HOM vs WT DEG’s per mutant genotype. (C.i-C.iii) The top twenty most up and down regulated genes in homozygous mutants versus control (D.ii-D.iii) HOM vs WT mutants effect size estimation of differential immediate early gene transcription. padj < .05, \**p <* 0.05, \*\**p <* 0.01, \*\*\**p <* 0.001. (F.i-.iii) Selective and possibly explanatory GO terms from HOM vs WT DEGs.

Pathway analysis of dysregulated genes was particularly remarkable for *kcnd2* and *wnt8b* mutants (Figure 4Eii-iii), with many Gene Ontology (GO) terms involved in axon development and synapse formation dysregulated, as has been reported in the context of human epileptic brain tissue[27]. *Arfgef1* mutants showed enrichment for genes related to neurotransmitter transport (Figure 4Ei).

## Discussion

As genomic sequencing continues in research studies and in the clinical setting[28], new potential candidate genes are being identified on an on-going basis. For epilepsy, as clinical sequencing guidelines advocate for a genome-wide approach, there are numerous possible findings each time a patient is tested and an ever-growing number of candidate genes emerging for human epilepsy [11]. There is thus a growing need for experimental models to provide evidence to support the role of these candidate genes in human disease, a first step toward precision diagnosis and treatment. Knowing the potential of the zebrafish system to model key epilepsy-related features, we focused on a subset of candidate epilepsy genes that are amenable to study in the zebrafish system. We evaluated 48 zebrafish models for epilepsy-related phenotypes— in local field potential recordings and structural abnormalities associated with hyperexcitability—and provide strongest evidence supporting the role of the human orthologs of *arfgef1, kcnd2*, and *wnt8b* in human epilepsy. *ARFGEF1’s* human disease association has been linked to developmental delay, variable brain malformations, and early onset refractory epilepsy[29]. Since initial generation of our candidate gene list and models, *Arfgef1* mouse models have been shown to have decreased latency to tonic-clonic seizures in both a PTZ-provoked and electroconvulsant threshold models[30], adding further evidence in support of this gene’s role in human epilepsy. Human alleles in *KCND2*, with predicted gain and loss-of-function, have been associated with global developmental delay in one report[31] and with tonic-clonic seizures in a pair of monozygotic twins in another [32]. *WNT8B’s* location at human chromosome 10q24 has raised the possibility of association with disorders including glioblastoma multiforme, spinocerebellar ataxia, and focal epilepsy[33]. *Kcnv1* and *ubr5* models showed some evidence of seizure-like movement, but given the permissiveness of the behavioral screen, additional study will need to be conducted.

Our approach was to prioritize and model genes, leveraging human population data (from affected and control populations) followed by behavioral swim pattern assays to identify larval zebrafish models with readily detectable seizure events. The zebrafish SII behavioral seizure event is comparable to the tonic and/or clonic seizures observed in many patients with epilepsy and provides an objective, quantifiable measure that can be assayed in initial studies to implicate genes in epilepsy, as we have done, and in pre-clinical compound screening studies. The 5 models showing seizure events were then assayed using conventional, albeit lower throughput, electrophysiology (LFP) recording to assess for evidence of hyperexcitability within the CNS. This approximates the electroencephalogram (EEG) conducted in individuals with suspected seizures as part of their epilepsy evaluation. Additionally, we demonstrated features pointing to potential mechanisms involved in generating or maintaining hyperexcitable brain networks, including decreased interneuron number in some mutant lines. Further, when we evaluated for convergent mechanisms using mRNA expression studies, we identified abnormal immediate early gene expression and dysregulation in genes related to axon development, synapse formation, and neurotransmitter transport.

The ease of CRISPR sgRNA guide design, generation, and delivery into 1-cell zebrafish zygotes allows for the evaluation of many genetic models efficiently. We had initially sought to model 81 candidate epilepsy genes that are intolerant to variation in the human population and that share good homology between human and zebrafish. We achieved 48 viable models, leaving the possibility that some of the other 33 may be disease-associated but that loss of function in zebrafish is not compatible with survival. Our method of assaying behavior as an initial screening tool affords efficiency but has limitations. Some zebrafish models such as *scn1lab*[17] have been shown to have frequent SII events and a robust hyperexcitable LFP pattern. Given the relatively small numbers of seizure events in the genetic models we generated, large numbers of larvae were required to conduct blinded experiments, and it are likely false negatives among the lines we designated as not having seizure events. We employed a binomial statistical method to compare populations of larvae (WT, HET, HOM for each gene) since this method is suitable in the setting of rare events. However, some models may have infrequent seizures, which we may have been missed given our relatively short sampling periods. A previous report showed mutant *arx* larvae have hypoactive movement with seizure-like activity via LFP[21], suggesting that low swim velocities and lack of SII events should not exclude the possibility of a gene’s causative role in epileptogenesis.

While the experimental design of this CRISPR-mediated reverse genetic screen was intended to create out-of-frame indels, similar to a previously described zebrafish screen of established catastrophic epilepsy genes[21], it did not guarantee generation of KO models. Interestingly, there is a growing body of evidence for increased transcription of compensatory genes in KO zebrafish models[34-36]. The mechanism for zebrafish KO compensation is still not completely understood, but nonsense-mediated decay (NMD) initiation from premature terminating codons (PTCs) has been shown to be necessary for an increase in adaptive gene transcription[37]. Of the 48 viable mutant lines 40 had predicted truncating mutations (Supplemental table 2), while three—included among the five genes (*arfgef1, kcnd2, wnt8b*) for which we demonstrate evidence for hyperexcitability—did not, in fact, contain truncating mutations but rather in-frame deletions. Interestingly, these three in-frame mutants proved to have the most striking evidence for epilepsy-like activity in either primary seizure detection or downstream assays. Additionally, it has been shown that far less transcriptional compensation occurs in HET PTC zebrafish models compared to HOM[36]. This phenomenon could potentially explain the discrepancy between significant SII seizure event scores in the motion tracking paradigms of HET truncating alleles (Supplemental Table 3) and the absence of corresponding findings in the corresponding HOM models. Three of five mutants with seizure events detected (*kcnd2, ubr5, wnt8b*) had seizures solely in HET larvae, including two with the most striking findings in downstream assays. The rate of SII seizure detection (5/48) and hyperexcitability on LFP (2/48) among our candidate epilepsy gene models is not surprisingly slightly lower than the rate (8/40) in a recent publication characterizing established epilepsy genes in the zebrafish system[21].

Even considering the complexities of zebrafish compensatory mechanisms and the methodological limitations of primary F_2_ behavioral screening, our pipeline does yield evidence supporting an association with epilepsy for 5 genes, with hyperexcitability on LFP adding stronger evidence for 3 of those 5 genes. *Arfgef1*[30, 38] and *wnt8b*[39] are associated with neurodevelopmental processes, and their models showed a reduction in inhibitory interneurons, while *kcnd2* encodes potassium channel subunit and its model had the strongest evidence of hyperexcitability via LFP recording. The approach we have employed, primary evaluation of behavioral seizures and abnormal electrical activity as well as structural and transcriptional abnormalities, attempts to accommodate the expected heterogeneity of the genes involved in human epilepsies by focusing on key core features of the network dysfunction common to all epilepsies. The models we have generated may now serve as pre-clinical models for drug screening, and the methods employed here can be applied to other emerging candidate genes for epilepsy.

## STAR★Methods

### Generating a candidate epilepsy gene list

Genes possibly associated with epilepsy were identified from the literature, using PubMed keywords “epilepsy,” “seizure,” and “gene” and published data from the Epi4K Consortium5 and other databases, including EpiGad (https://www.epigad.org/), Carpedb (http://carpedb.ua.edu/), and OMIM (https://www.omim.org/). Gene lists obtained from a compendium of clinical genetic labs (https://www.ncbi.nlm.nih.gov/gtr/) were also included. After removing duplicates, a list of unique “candidate epilepsy genes” was consolidated and evaluated by the neurologists and genetic counselors of the Boston Children’s Hospital (BCH) Epilepsy Genetics Program.

### CRISPR guide design

We used the CRISPR/Cas9 system[40] to disrupt the function of the zebrafish gene paralogs of the 81 candidate epilepsy genes described above. CRISPR/Cas9 sgRNA guides were designed to introduce out-of-frame deletions predicted to result in premature terminating codons (PTC) of target genes via non homologous end joining. Two sets of CRISPR/Cas9 guides were designed near the 5’ and 3’ ends for each transcript, excluding the first exon and last exon, respectively. Designs for duplicated genes included two sets of guides per paralog (e.g., *atp1a3a* and *atp1a3b*). Target sites were selected on different exons across the entire length of each non-overlapping gene transcript using CHOPCHOP[41] (http://chopchop.rc.fas.harvard.edu). Guide requirements included targeting efficiency >50% based upon high GC content and minimal self-complementary within the sgRNA, or between the sgRNA and RNA backbone. Additional requirements included no-off target matches and a Protospacer Adjacent Motif (PAM) sequence (nucleotides NGG) upstream of the desired target (Supplemental Table 1). All guides were generously donated by Synthego (Redwood City, CA, USA).

### Generation of stable zebrafish mutants

Zebrafish embryos (background strain TAB) were injected at the 1-cell stage with 500 pg/nl sgRNA and 500 pg/nl Cas9 protein (Invitrogen, Waltham, MA, USA). Mosaic F0 larvae were raised to breeding age (2-3 months) in 2.8L tanks containing fewer than 30 zebrafish and outcrossed to a WT casper transparent line. Germline transmission of CRISPR indels was assessed via genotyping pools of 5 2 dpf F1 TAB/casper larvae from isolated F0 parents, while remaining F1 clutches were housed pending genotyping results. Mutant allele-containing F1 fish were raised to breeding age and fin-clipped for DNA to assess for HET mutants. CRISPants were generated using Cas9 protein and synthetic guides. Genotyping of individual or pooled larvae was conducted by Sanger sequencing from DNA extracted by standard methods. F1 HET fish were in-crossed to generate F2 larvae for additional assays. Mutants cnnm2a and cnnm2b were the only exception to this strategy and were imported into the lab as stable HET stable lines (generous donation from Dr. Summer Thyme, University of Massachusetts Chan Medical School). A subset of stable F2 mutants were crossed into transgenic GCaMP6s and tg(*dlx6a-1*.*4kbdlx5a/dlx6a: GFP::vglut2:DsRed*) (generous donation from Dr. Ellen Hoffman, Yale University). HET F3 transgenic fish were sorted for maximum fluorescence and genotyped.

Conditions: F2 larvae were kept in 100ml Petri dishes in sterile fish water (SFW) with 2% methylene blue at a density of fewer than 100 embryos per plate. Methylene blue SFW was cleared at 1 dpf and replaced with 100% SFW before larval chorion shedding[42]. Larvae were raised to 5 dpf with daily replacement of SFW and plated into 96-well plates with 100 ul SFW per well.

### Identification of seizure events in larval zebrafish

F2 5dpf larvae were placed in an automated motion tracking system (DanioVision, Noldus, Wageningen, Netherlands), allowed to acclimate for 5 minutes and then evaluated in three conditions: (a) one hour of recording spontaneous activity, (b) a 26-minute light paradigm consisting of four oscillating periods of dark and flashing light (2hz-10hz with a 2hz increase per light interval, a paradigm adapted from human EEG methods to detect photosensitivity[43], and (c) 30 minutes in subthreshold PTZ paradigm (adding 100 ul per well PTZ well for a final concentration of 0.125 mM). Subthreshold concentration was determined by the highest PTZ concentration at which no SII events were observed in WT larvae during a 30-minute PTZ titration trial. Motion detection assays were conducted in duplicate for each gene. Assayed larvae were generated from HET x HET in-crosses to provide internal WT controls and assess for batch effects. After screening ∼20,000 larvae, a list of the most promising candidates for a potential seizure model was developed. Criteria were developed based on SII stereotypical seizure-like swim patterns found in PTZ-treated larvae, as described[17].

Motion tracking data were analyzed using Ethovision Xt software (Noldus, Wageningen, Netherlands). SI seizure events were defined by significant increase in the Ethovision parameters “average velocity,” “average distance” compared to WT controls. SII seizures were identified by auto-detection of “total distance” (TD) >60mm and “velocity max” (VM) >90mm/sec, evaluating data from 5-second bins; seizure events were confirmed by visual inspection by a trained researcher. We calculated the binomial probability of the proportion of mutant larvae displaying SII events compared to proportion of the number of WT displaying SII events (Supplemental Table 2). Initial population probability of spontaneous WT SII behavior was determined (0.023) from ∼400 uninjected TAB larvae. Population probability was further refined after incorporating conspecific WTs from a subsequent series of genotyping performed on any gene with significantly higher binomial probability of larvae with > 0 SII events given suspected Medellin inheritance with the WT population proportion of 0.023. From this cohort, a new proportion of WT > 0 SII event was updated to a frequency of 0.027. The updated conspecific WT genotype SII swim data was then incorporated into the population probability for binomial significance assessment in the PTZ and light-provoked paradigms to be use as the threshold for behavior screening. These evaluations were conducted blind to genotype, and post hoc, larvae were sacrificed by standard freezing methods and genotyped via Sanger sequencing.

### Electrophysiology: local field potential recording and analysis

Electrophysiological studies were conducted using established published methods[17]. 5-6 dpf larval zebrafish were treated with 2 mg/ml alpha-bungarotoxin (Invitrogen, Waltham, USA) for ten minutes and embedded in 2% low melting point (LMP) agarose (Fisher Scientific, Cambridge, MA, USA) to eliminate movement. Embedded fish were perfused with artificial CSF throughout the recording. A glass microelectrode was placed in the optic tectum, the largest forebrain region in the zebrafish, with CSF containing 117 mM NaCl, 4.7 mM KCl, 2.5 mM NaHCO3, 2.5 mM CaCl2, 1.2 mM MgCl2, 1.2 mM NaH2PO4, 11 mM glucose (Sigma-Aldrich, Natick, MA, USA). Electrodes (1-7 MΩ) were backloaded with 2 M NaCl prior to placement. Electrical activity was recorded using pClamp software (Molecular Devices, Sunnyvale, CA, USA) and an Axopatch 200B amplifier (Axon Instruments, Union City, CA, USA). Voltage records were low-pass filtered at 1 kHz (3 dB; eight-pole Bessel), high-pass filtered at 0.1–0.2 Hz, digitized at 5–10 kHz using a Digidata 1440A interface. Proper electrode positioning and detection of physiological activity was detected with the addition of PTZ at the end of each recording and data corresponding to animals failing to respond to proconvulsant was disregarded.

Raw EEG data were normalized using a root mean square (RMS) method, band-pass filtered at 3-10Hz to detect events in the physiological 100-300msec range and then smoothed with a 200-msec boxcar moving average filter. For display, the spike amplitude was normalized, and the “instantaneous spike rate” was computed by summation of spikes in a 20-second moving window. Spikes were defined as deflections exceeding the interquartile range (IQR) of filtered voltage values by 2x within a 10-second moving window (1000 samples). The peak of a “spike” was defined as exceeding 6X the moving IQR. Spike counts between all genotypes ranged from >100 to ∼500 within recording window with median amplitudes from .05 - .4. Additionally, mutants displayed an increase in deflection clusters and durations. Burst were defined as greater than 3 spikes for longer than a second at a rate exceeding .3hz. Bursting event counts within all genotypes ranged between 0 – 30 within analysis window counts with durations ranging from 0-1000 sec.

### Imaging

Confocal imaging: F3 tg(*dlx6a-1*.*4kbdlx5a/dlx6a:GFP::vglut2:DsRed*) HET larvae were crossed and sorted to maximize fluorescence and pigmentation loss. 5 dpf larvae were fixed in 4% PFA overnight and stored in PBS at 4 degrees Celsius. Larvae were dorsally mounted in 100 ul 1.2 LMP agarose. Imaging was conducted on a Zeiss LSM700 at 10x magnification with a 10 um step size. Image analysis was conducted using Fiji software, as described previously[44]. Fluorescence intensity was normalized between samples with a cutoff threshold of 99% greater than background. GFP-expressing interneurons were quantified.

### RNA-Seq analysis

Three replicate larvae from Het in-crosses, aged 5 dpf, were sacrificed on ice and stored in RNAlater (ThermoFisher) according to product specifications. Larval heads were isolated from bodies, which were used for Sanger sequencing for genotyping. Heads were pooled in groups of 20-25 according to genotype and RNA was isolated using a Qiagen RNeasy plus kit according to specifications. QC was performed via Agilent 2100 bioanalyzer, and only samples with a RNA integrity number value of >5.8 were used for library preparation. Sequencing was performed using an Illumina NovaSeq 6000 Sequencing System with >20 million reads per sample. Sequencing data were aligned to zebrafish genome danRer11 using STAR. Expression was quantified using featureCounts. For each group of HET, HOM and conspecific WT samples, only protein-coding genes with transcript per million value >= 1 in at least 2 samples were retained for downstream analysis. RUVSeq (RUVr k=2, v1.28.0) was used to remove batch effect; differential gene expression analysis was then performed using DESeq2 (v2.28.0). Differentially expressed genes were defined as those with absolute log2 fold change > 1 and adjust Pvalue < 0.05. Gene Ontology (GO) enrichment analysis was performed using clusterProfiler (v4.7.0).

## Acknowledgments

The authors are grateful for the support and encouragement of the family of our dear friend and colleague Jeremy F.P. Ullmann, PhD, who began the work presented and died in a tragic hiking accident in 2019. We thank the physicians and genetic counselors of the Boston Children’s Hospital Epilepsy Genetics Program for insights into the human genes to consider modeling, the dedicated leadership and staff of the BCH Zebrafish Aquatic Facility, and the Rotenberg Lab and the IDDRC Animal Behavior and Physiology Core at Boston Children’s Hospital, funded by NIH/NICHD P50 HD105351. We thank Drs. Summer Thyme (University of Alabama) and Ellen Hoffman (Yale University) for valuable genetic zebrafish lines. CRISPR sgRNA guides were provided by Synthego.

Funding for this project included support from the BCH Translational Research Program, the 2021 Boston Investment Conference (AP), and the National Institute of Neurological Disorders and Stroke K08NS118107-01 (CMM).

---

**Supplemental Figure 1.**
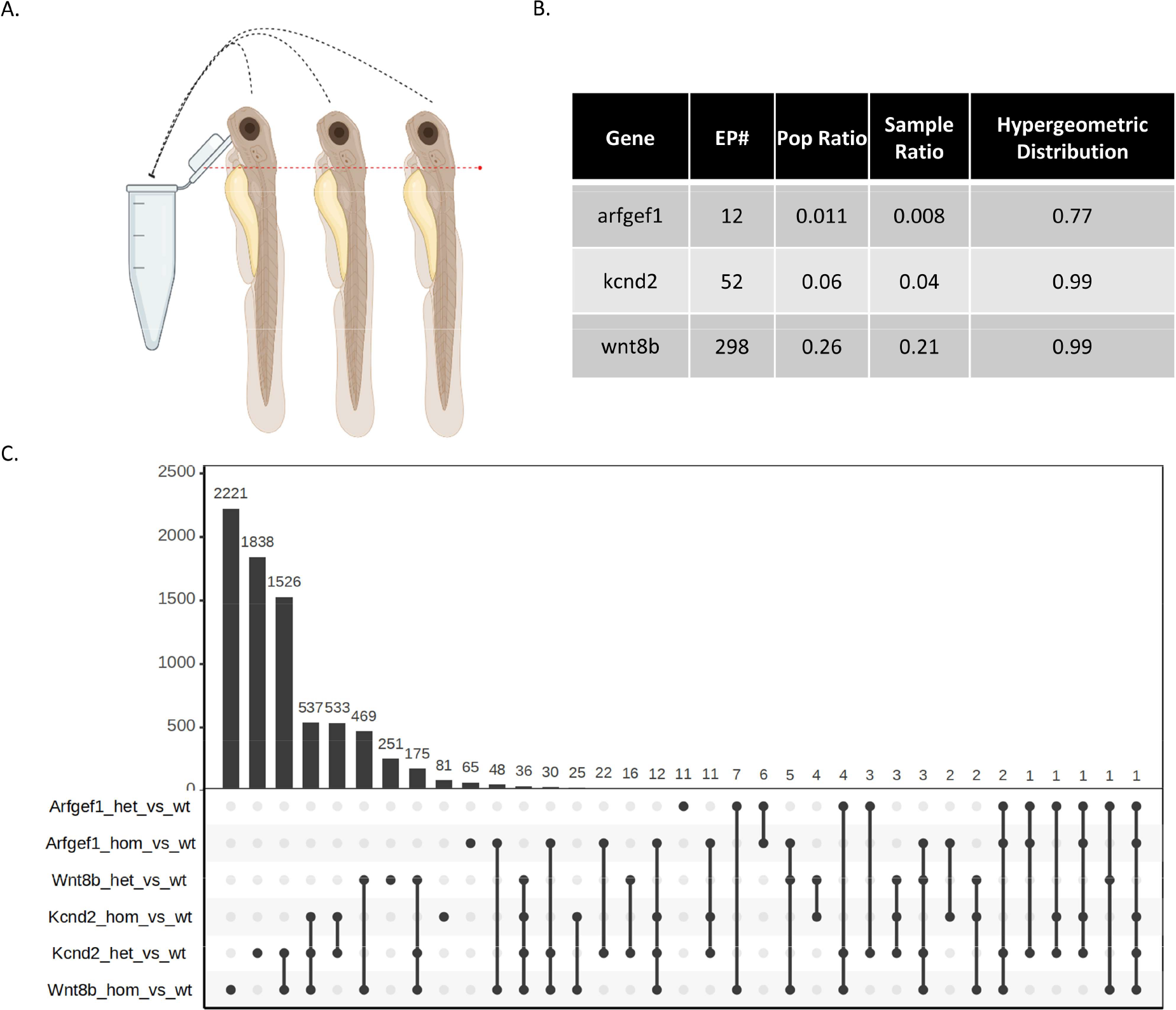
(A) Schematic representation of RNAseq assay and head isolation. (B) Co-occurrence of mutant HOM DEG’s with EPGP gene list (EP#) and significance via the hypergeometric distribution based on population ratio and sample ratio occurrences. (C) DEG co-occurrence numbers between arfgef1, kcnd2, and wnt8b mutants.

## Notes

### Competing Interest Statement

The authors have declared no competing interest.

