## Supplemental Table 1 for "Zebrafish models of candidate human epilepsy-associated genes provide evidence of hyperexcitability"

| *Homo Sapien* Gene | *NCBI Ref Seq* | GnomAD Constraints pLI | Missense Z | *Danio rerio* Gene | NCBI Ref Seq | Percent Identity (%) | Transcript Variant | Transcript Target | CRISPR Guide | Fwd Primer Sequence | Rev Primer Sequence |
|---|---|---|---|---|---|---|---|---|---|---|---|
| *ABCC9* | NP_005682.2 | 0.48 | 4.97 | *abcc9* | XP_005164758.1 | 79.26 | 201 | 5' | GGTTTCCTCCATCTTGGAGGCGG | TTACACACCCTCAAGTGGTTCC | TACCAACCAATAGGCGAGGC |
| | | | | | | | | 3' | GTCGTTCCCACAGAAAGACAGGG | GCTTCTTTAGCCCAGTCAGG | CAGAATGAAACGCAACTCTGTG |
| *ACO2* | NP_001089.1 | 0.42 | 2.92 | *aco2* | NP_944590.1 | 82.99 | 001 | 5' | CCGCCGCCACATGAAAACGGCGG | CGGTCGATTGAGTCTGCAAC | TGCACAATAAACATGAGCTCCC |
| | | | | | | | | 3' | AATAGCACTTACCTTTCCAGGGG | GCAATGTTTAGAAACGCGCTC | GAGAACATGCTGCTCTGGAG |
| *AGPAT3* | NP_001032642.1 | 0.14 | 2.54 | *agpat3* | NP_998590.1 | 75.33 | 001 | 5' | GTCGAAGGGCCACAGCACGCAGG | | |
| | | | | | | | | 3' | GTCGAAGGGCCACAGCACGCAGG | TTAGGACCCACACAGACCAC | CTTCCAGCAGTGTGAGTGTG |
| *ARFGEF1* | NP_006412.2 | 0.14 | 5.37 | *arfgef1* | XP_009295669.1 | 85.11 | 001 | 5' | CAGGGCCCGCAGACAGACGAAGG | TCAATTCTCGTGGTGAGCAGG | TTGGTGAGCTTGACCGGTTG |
| | | | | | | | | 3' | CAGCGTTTGCAGTGACGCAGTGG | AGCAGAGATGCTTAGCCAGG | AATACTGCTGTGACAATGACGG |
| | | | | | | | 002 | 5' | TCCAAACTCTCTGGAAGAGGAGG | GTGCGTTCACATATTAGCAGG | GTTACAGAAATGAAGGCAAGCC |
| | | | | | | | | 3' | TACACTCTGTGCATCTGCGGGGG | TTGCTCCACTCCACCATACAC | TGTGTAGACAACAGTCTCTGGAC |
| *ATP1A3* | NP_689509.1 | 0.06 | 6.33 | *atp1a3a* | NP_571759.2 | 90.67 | 001 | 5' | GTGTTGACGTGTCTTACCAGTGG | GAGATCCACCCACTGCAAAC | ATGACCTTAGCTGACAAGAGGG |
| | | | | | | | | 3' | CCTGCTGGAACACAGAGTTGCGG | CACTGCTCAATGTGAATGTGC | CCCCTAAGACTAACAATGTGCAC |
| | | | | *atp1a3b* | NP_571760.2 | 92.17 | 001 | 5' | AGGAAGAACACGTACCACCAGGG | TTCTGTCTGGCCTAACTCATGAC | TCCTCAAATGGCTCTGCAGC |
| | | | | | | | | 3' | GCCAGTGTAATACTTACAAGGGG | GCTGTGGTCATTATTCCTGAAC | TGAGGCACACTGATCATTCTC |
| *CACNG2* | NP_006069.1 | 0.38 | 2.09 | *cacng2a* | NP_956935.1 | 88.65 | 001 | 5' | GTACATATCAGCCAATGCGGGGG | CCCCTGATGAGGTATGAACAGG | AATCTCGGTGGAAGGTAGAGC |
| | | | | | | | | 3' | TCAGATGTTATTGACCACGGTGG | AGGACAAACGATTCGGAAGGG | CGCCTTACCTTCCAAGCAAC |
| | | | | *cacng2b* | NP_001186659.1 | 82.98 | 201 | 5' | AGACTTGTAGAACTCGCTGGCGG | GTGTGTGTGTGTGTGTGCAG | AGCTCCTCCTCAAATGCCTG |
| | | | | | | | | 3' | GTATGGGGGTGTTTGAGCGCGGG | TGATGCCAGAATGCTCCAATTG | ACATCCGAAACATGCAGTGC |
| *CACNG3* | XP_693299.1 | 0.20 | 2.64 | *cacng3b* | NP_006530.1 | 80.06 | 001 | 5' | CATGGGCGAAGCGTCACGAGAGG | TCTCGCCCTCTTCTCTTCAC | TGTTGGCCGCATCTTTGAAG |
| | | | | | | | | 3' | GATAGTCATCAGGCTGAACGCGG | GTGACAGTTTTTCCACAGCCC | AGCACATGAGACGCCTTACC |
| *CALM2* | NP_001734.1 | 0.37 | 2.79 | *calm2a* | NP_956290.1 | 100 | 001 | 5' | GCCGGTAAGACATTGGCACGTGG | AGGTTTTCCAGTAGTCAGTTCC | GGGGAAAACCATGGGAATCC |
| | | | | | | | | 3' | GAATGGATACATCAGCGCAGCGG | TCACCTTCGTAGTTAACCTGTCC | AGGTACTGTCATGCGCTCAC |
| *CALM3* | NP_005175.2 | 0.36 | 2.98 | *calm3b* | NP_955864.1 | 100 | 201 | 5' | TCTACTCCTTCCTCAGGCAATGG | | |
| | | | | | | | | 3' | AAGAATTAGGGACTGTCATGCGG | CTAGAATTCAAGGAGGCCTTTTCC | GCCGTCACCATCAATATCTGC |
| *CHD2* | NP_001262.3 | 0.07 | 5.21 | *chd2* | XP_021323365.1 | 70.43 | 202 | 5' | CGTGGGTCAATCAGAGGCGGGGG | TCTCCTCGTTCCAAAATGGCAG | AAAGAGAAGGCGCTCGGATG |
| | | | | | | | | 3' | CAGGAAGAAAGACAACCTGGCGG | GGCATTCTGAATCCAACAGC | AAAGAGTGTGATAGTTCAGGGG |
| *CHD4* | NP_001264.2 | 0.16 | 6.34 | *chd4a* | XP_021323760.1 | 78.21 | 001 | 5' | TGCAGCCTCACTGCAACAGGCGG | AGCTCTTGAGTTCTCGTAGACG | GGGGGCGAATAATTCTGACTTC |
| | | | | | | | | 3' | AAAAGTAGCCGCGAGAGCAAAGG | TCGTACTTGTGCAGATGTCCG | TATCCACACACACACTCGCAG |
| | | | | | | | 002 | 5' | GTGTGTTTGATCCACAGAAGAGG | | |
| | | | | | | | | 3' | GTCTCTATCGTCCTCACTGCCGG | AAAAACCCGGCAACACAAAGC | GTTCTTTTATTTTCTGCCTCCCCC |
| | | | | *chd4b* | XP_021328683.1 | 77.5 | 001 | 5' | TAAGGAACCAGAGCCAGAGGAGG | CAAAACTGTAGCTGGTGAAAGC | TCCTTTTGGTGGATTCGTGG |
| | | | | | | | | 3' | ACCCGAGGTCACGCAGACGCAGG | ATGCGGACCTGTTCAAACATC | ACGCATCTGATCACCTGAGC |
| *CLCN4* | NP_001821.2 | 0.15 | 4.52 | *clcn4* | NP_001070786.1 | 90.19 | 201 | 5' | TATCGTCCCTCTGATGGCGGCGG | GTGTGTGTGTCATACTGATAGCC | CTGGTTCATTCTGCTGTGGC |
| | | | | | | | | 3' | ACTGAGGAGATGAACGGCGCCGG | ACAACTGGGATCGACAACCC | TGTCCAGTATCATGCAGCCC |
| *CNNM2* | NP_060119.3 | 0.27 | 4.41 | *cnnm2a* | NP_001138257.1 | 76.86 | 001 | 5' | GAAGGAGAAGCATTACGCGAAGG | AAAGTGATCGTGGTGGAGGAG | ACAGTTCCAATCTCCTGCCC |
| | | | | | | | | 3' | GAGGCCGTGACTCCAACACTGGG | ATGCGACATCAAAGACGAGC | TGTGCAAACTGAACCATGAGC |
| | | | | *cnnm2b* | XP_009290541.1 | 79.52 | 001 | 5' | CGAGGGGGTACTGGAACCGACGG | ATCAGTAGAGGCACGAGTTCG | CTGTCGATGTTGCCGATCAC |
| | | | | | | | | 3' | TGTCTGGTTGAACAGCGTGGTGG | TGATCCACAGACTGGTACCG | AAGCGGGTCACTGGACTATTC |
| *CSNK1G2* | XP_005259555.1 | 0.25 | 2.80 | *csnk1g2a* | XP_005171263.1 | 76.6 | 001 | 5' | GGAAGAGGAAAGCGCTCCAGCGG | CTTGCATTTGCTCTGGGTGG | AGTGGATGCCGAAAGTGAGG |
| | | | | | | | | 3' | GGTTGGCCCGAATTTTCGGGTGG | AGGAAAAGGAGAAGCGGAGG | GTTCATGAGATCGGCAAGTACC |
| | | | | *csnk1g2b* | NP_001039315.1 | 75.78 | 002 | 5' | GCACCTGAAAGTAAATACAGAGG | TGTTCCTCGTTGTGCCTCAG | CAAGTTCTTTTCCACGAGCGC |
| *CUL2* | NP_003582.2 | 0.23 | 4.75 | *cul2* | XP_002666627.1 | 92.21 | 001 | 5' | ACACCATGTCCTTAAAGCCGCGG | CATTGCACATCTCCAAACCCG | AGGCCTATCTAAAGGGTTATGCTG |
| | | | | | | | | 3' | GGAACAGACGAGGAGCGCCGTGG | TAACAGTGGTGTGTGCTGC | AGCCTGAATCCACTTTTTCCC |
| *DIP2C* | XP_009295792.1 | 0.15 | 4.67 | *dip2ca* | NP_055789.1 | 87.96 | 001 | 5' | ACGGTGGTATCGTGAAGCGGAGG | TCTCTAAGGCCGTGTGTGTG | AAGAATCCTGAGGGACCAGTG |
| | | | | | | | | 3' | ACTCTCGTGGAGAGAAACAGAGG | GACTCTGTGTGTTCCAGTGC | AAAAGATGTCGCAACACCTCAC |
| | | | | *dip2cb* | XP_021326954.1 | 76.94 | 001 | 5' | CTCAGATGTTCACTCAGAAGCGG | AGCAGTACACACACACCTGG | TCTGTGGTTAGCACTGTCGC |
| | | | | | | | | 3' | ATGGACGAACCTCTTGGTGGTGG | GCGTTGCTTTTCTCCACGAG | AGGTGATGAGGGGTGCAATAC |
| *EFTUD2* | NP_004238.3 | 0.09 | 4.03 | *eftud2* | NP_956802.2 | 91.06 | 201 | 5' | TCTACGGGACACAGAGACGGGGG | | |
| | | | | | | | | 3' | AGAAGTGTATGGTCCTGAGGTGG | GCACACACACACTCACACTG | ACACACATGGAAGAGAGGTGG |
| | | | | | | | 202 | 5' | CTTAAAACAGGCGGATGAGGAGG | TCCATGTCATAGACGGTCGC | TCAAGCCGGGAATCGAACTC |
| | | | | | | | | 3' | TGGCGAGCTCCAACAACATGGGG | ACTTGCCCGTTCTTCACCTC | GTGATGTTGAGCTGCGGAAG |
| *FAM50A* | NP_004690.1 | 0.29 | 3.02 | *fam50a* | NP_001017636.1 | 86.22 | 001 | 5' | AAGGAGCGGCCAGTGAAGCGGGG | | |
| | | | | | | | | 3' | CGGGATCGTAAGGTTCCCATCGG | TTGGAATGCGGTGAACTCG | ATTTGTAACGTTTGCCTGCC |
| | | | | | | | 003 | 5' | ACTAGGAAAGAACCCAGACGTGG | ACCGAGTCATGCGACAATGG | TCTGCCGACATCCTGTTCAC |
| | | | | | | | | 3' | CGGGATCGTAAGGTTCCCATCGG | ACCTCTGTGTGTGTTTGTGTG | ATTGCATCCTGCCATTTGGG |
| *FAR1* | NP_115604.1 | 0.30 | 3.30 | *far1* | XP_005174372.1 | 77.61 | 001 | 5' | GTGTGCATAGCAGTTAGGAGAGG | CCTGTCCTATCAATGCCAAGTTG | GTGCTTCATGGATTGGAACGG |
| | | | | | | | | 3' | CACTTTCCCCATAAATCCCGTGG | CCGGTGCAGTAATACAACACC | ACAGTTGATCATGTCAGCGATG |
| | | | | | | | 003 | 5' | TGGAGCGCAGGTTAACATTGGGG | TGGATACTGGGTTCTGCACG | TCAGTCAGTCATGCCACACC |
| | | | | | | | | 3' | TTTTCCCACCCCCTAGGCTGCGG | | |
| | | | | | | | 004 | 5' | CAAGAGGAACCCCCTCGAGCAGG | AACAGTTCTCGCGTTTCTGC | ACACACACACAAACACGCAC |
| | | | | | | | | 3' | TTTTCCCACCCCCTAGGCTGCGG | | |
| *FGD1* | NP_004454.2 | 0.20 | 3.52 | *fgd1* | XP_688794.4 | 75.45 | 001 | 5' | AATCACGTGCCACCATGCCGAGG | GTGATTGTGCACCAGTCTCAC | CGGGGTGATTTGTTCTGCAG |
| | | | | | | | | 3' | GTTTTATCCCGAGCCGGCAGGGG | AAACAAAGCCTTGCAGCTCG | CATTTGTACAGTTTGCGTGTGC |
| | | | | | | | 201 | 5' | GATTCCCCCTGATCAGCCGGCGG | AAAGGTCATTGCGCATTGGG | GCACATAGGATGCAAGCTGC |
| | | | | | | | | 3' | ACAGAAGCAGGCCTCGGTGGCGG | GTTCATTCCGCTGTGGTGAC | TCTTCTCACCTGTGGAGCAC |
| *FLNA* | NP_001104026.1 | 0.08 | 3.78 | *flna* | XP_009295045.1 | 64.83 | 004 | 5' | AGCTATGTGCCCACAAGAGAGGG | CACTGCAGGTCAGAGTTCAG | GGGTGAAGTGGTATCTCTATGG |
| | | | | | | | | 3' | TAGCCTGTAAAGAAACACAAGGG | TTGCCCTGCTAGCTCGTTAG | TTGGCCACTTTTCCAGAGGG |
| | | | | | | | 201 | 5' | AGCGTTGGATAGGGGAGCGGCGG | CACCCAAAAAGCGAGACAGC | AGACGTTCTCCAGCTGCATC |

| | | | | | | | | | | | |
|---|---|---|---|---|---|---|---|---|---|---|---|
| | | | | | | | 201 | 3' | GGTAGGAAACGTTGTAGAGGCGG | TTGGCTTCACAAGGGAGAGC | GATGGATGTTTTTGTGGAGGGC |
| *FLNC* | NP_001449.3 | 0.25 | 2.79 | *flnca* | XP_002666982.1 | 78.72 | 001 | 5' | TGAGAATGTCTGCTTACCAGGGG | GCAAAGCCATTGTAGACGGG | TCACCTAATCACAACCGCCC |
| | | | | | | | | 3' | GGGGGGACGAAACTATACCAGGG | CATGCTCATGGTTGGTGTCC | CCCATCACCATCAAAAGTGCC |
| | | | | *flncb* | XP_009298480.1 | 82.28 | 001 | 5' | TACACGAAGGGAGCAGGCAGTGG | TGCTCATTCTTTTGCTGGGC | ACCCCAAGTAACGGTGATGATG |
| | | | | | | | | 3' | TCAGGTGCACGTCTGTCTGGCGG | TGTGCCACAGTTTGCCAAAG | ATGAAGGCTTTGGAGAGGCC |
| | | | | | | | 201 | 5' | AGTTATCCACCAATGCACCGAGG | CAGAAAGCTGACGCCCAAAC | TTATCCCCCAGCCCTTGTTG |
| | | | | | | | | 3' | TCAGGTGCACGTCTGTCTGGCGG | | |
| *FRMD4A* | NP_060497.3 | 0.14 | 2.46 | *frmd4a* | XP_009291718.1 | 79.38 | 001 | 5' | TTAGCCCAAACTAATGGCGAAGG | ATCTGGGCATTCTTGAAGGG | CCCATGAACATTGCTTGAGC |
| | | | | | | | | 3' | GTTTGGAGAACGACCAGGAGGGG | GAACTTGGCAGCCCGAAATG | GCGGCTGTGCTCTTTAATGG |
| *GABBR2* | NP_005449.5 | 0.17 | 4.63 | *gabbr2* | NP_001137515.1 | 78.83 | 001 | 5' | GTAGCAGATTGCATCAAAGAAGG | AGCGCTGCAGCCTATATAAAGAG | TACGTTTCTGGCCCGACTTC |
| | | | | | | | | 3' | GATCCTTCTAGGATAAGTAGAGG | TGACATCCTGGTGGAATGGC | AACTATGCAGGACAATGGCCC |
| *GLUD1* | NP_005262.1 | 0.61 | 3.06 | *glud1a* | NP_997741.1 | 83.87 | 001 | 5' | AGCGGAGTCTACGCACACCGAGG | GCCGTGGTGTCCGTTTATTC | ACTTGCCTCCTTTACACGGG |
| | | | | | | | | 3' | TGATTGGCTTGTCTCATAGGGGG | GGATTTCTGTGAGTGCCAAC | CAAGCATACGTTTCACCTAAGC |
| *HCFC1* | NP_005325.2 | 0.06 | 5.63 | *hcfc1a* | XP_005169361.1 | 77.15 | 001 | 5' | CGGTACCTCGACCCAGACACGGG | GTCGCTCACAGAAACAACTTG | ACGTTCTTTGGTTTGTACGC |
| | | | | | | | | 3' | GTTTTGGAGGCTAAATGCAGAGG | GCCATGAAATAGCCATGACTTGC | AGTCAGAGATGGGTTAAATCCGG |
| | | | | *hcfc1b* | NP_001122009.1 | 64.73 | 001 | 5' | AAGCGCTGGTCAGAAGATGGCGG | TAGCTCCTGTTCAACGAAATCC | ATTCCTGTGTTGATGCTGGG |
| | | | | | | | | 3' | GGTTGAGCTCATTAAGAAGGCGG | CCAAACCTGCAGCAAAGAGG | GAATGGGGTGAGAAGTGAAAGC |
| *HECW2* | NP_001335697.1 | 0.26 | 3.31 | *hecw2a* | XP_021334623.1 | 76.87 | 001 | 5' | CTAGAGTTTACACTGTGCCGCGG | TTATTCCGTAGAGTCGCCCC | GGCGTTGCTAAGCTAACTCTG |
| | | | | | | | | 3' | CTAGCTGTACCGTGGGAGGGAGG | GCCAATTCAACATGTGTCTCTG | ATAGAGCATGGAGAACGAGGG |
| | | | | *hecw2b* | XP_003197660.3 | 80.5 | 001 | 5' | GTGAAGTGGCGTATCGAGCGCGG | CCATTGTCTTACAGAAGTGTGTGC | CACTGACTCAAATCTGTCAAGCAG |
| | | | | | | | | 3' | TTCTCCACACAGAAGCGCCGCGG | CAAAGCATCATGGGGCATATG | GGTCAAATTTAGACAGACAGGC |
| | | | | | | | 201 | 5' | GTTTGGGACTCGAAGAACCGCGG | ACTTCAGCTGGAGGCCTTTC | GCTGAGCAACTGAAGTACCATAC |
| | | | | | | | | 3' | TTCTCCACACAGAAGCGCCGCGG | | |
| *HK1* | NP_277035.2 | 0.33 | 3.23 | *hk1* | NP_998417.1 | 85.22 | 001 | 5' | GAGGGAACACCCCTAAGTGCCGG | AGGACATTGCAGCACAAACC | ACAGCTTGACTTCCTAGGCC |
| | | | | | | | | 3' | GCCAAACGGTCACTAGAGGGAGG | TCCACAACAGCAGCCATTCC | ACCCTTGAACGGCAGTGTAC |
| *HNRNPH1* | NP_001351158.1 | 0.11 | 4.09 | *hnrnph1* | NP_991247.1 | 69.68 | 001 | 5' | GTGACGAAGCGTCTTTGAGGGGG | CGTGTGTGAGCACTGTTCAG | AACCTTCTCCATCCGCCATG |
| | | | | | | | 001 | 3' | TTAGGCGGCAGTTACTACGGCGG | ACAGAAGTGATTTGTTCCCGC | GGAACAGAAAACTCTTGTGCTCC |
| *HUWE1* | NP_113584.3 | 0.06 | 8.87 | *huwe1* | NP_001352583.1 | 76.35 | 001 | 5' | TGTTGCAGGCGGGCAAGCAGTGG | GCAACCCCAGACCAACAAAAC | AAGGCCTTTGATTTGCAGCTG |
| | | | | | | | | 3' | ATAGGTGACCCGATCGCCGGGGG | AGATAACGGCAACAGCAGGG | TAACGCTCTTGCCCAGGATG |
| *IQSEC1* | NP_001127854.1 | 0.22 | 2.28 | *iqsec1a* | XP_017209231.1 | 72.49 | 201 | 5' | GCGCAAGCAAGCAGAAGAAGAGG | GACTGTTGAAAGCAGTGAGCC | GGTGTTTCGTGACCCTTTAAAGTG |
| | | | | | | | | 3' | GAGACGGGGAACGGCAGCATGGG | GTACTTTAATAGGGGAGTGGTTCC | TGTTAAATCTGGACATCAGCGG |
| | | | | *iqsec1b* | XP_005169390.2 | 76.85 | 001 | 5' | AGTTCAAAGAACGTAGACGGAGG | GCGGTCACTCAACTCAACAG | ACGCCATTACAGAACTGGAGG |
| | | | | | | | | 3' | TACGTGGATCCCCAAAAAGGAGG | TCTCCGGACACAAACGCTAC | TCGGATTCGGTCACAACGAG |
| | | | | | | | 201 | 5' | AGTTCAAAGAACGTAGACGGAGG | | |
| | | | | | | | | 3' | TACGTGGATCCCCAAAAAGGAGG | | |
| *KCND2* | NP_036413.1 | 0.57 | 3.51 | *kcnd2* | NP_001076336.1 | 80.28 | 201 | 5' | TGGTTCCTAAGACCATCATGGGG | GGGTGAACTCATGAAACGAATG | GGACCTGCTTGTAGTTTGGG |
| | | | | | | | | 3' | CCCAATTCCAACATGGCCGGCGG | TACCATAAGTTCCCATCACCGC | TGTGTGTTTGTGTCAGTGTGC |
| *KCNJ2* | NP_000882.1 | 0.61 | 2.75 | *kcnj2a* | NP_001318145.1 | 83.72 | 001 | 5' | CCGAGGTTCTAAGGGAACGGTGG | CGTTGGACAACTGCACATGG | TGAAATCCTCTGGGCACACC |
| | | | | | | | | 3' | TAACGAACATCCATCGCCAGCGG | GAAGATGCGGTCAATGCCAC | TGAGGTGATGGGAAGTGTGC |
| | | | | *kcnj2b* | XP_021328818.1 | 71.27 | 001 | 5' | AAAGGACCGGCTCAAAGCGATGG | GTTGACATTCGCTGGCGTTG | GCTGCATTGAGGTGTGTTGG |
| | | | | | | | | 3' | ATATGGCTACCGCTATGTGACGG | | |
| *KCNJ3* | NP_002230.1 | 0.26 | 4.06 | *kcnj3a* | XP_002663356.2 | 83.1 | 001 | 5' | TAGTAGACCTTAAATGGAGGTGG | TAGTGACCACTACCTCCTCCAG | AGTTTCCGCCCTCTTCTTGG |
| | | | | | | | | 3' | AGCGCGGACATCATACACAGAGG | AACTGCTACCGCGTTTTATTCG | CTGTTCTTTTCTCCGCTGTTGC |
| | | | | *kcnj3b* | XP_005165913.1 | 75.96 | 001 | 5' | TGTTGCGCAGGTTCCCCACGCGG | TCCTAGGCTCTATAGTGGACGC | AGGTCCTTCTCATTTATCAGGGC |
| | | | | | | | | 3' | TGTTCTTTAACACTGTGCGGTGG | TGGCTCCTGTACTGGTAGATGG | GCTTCTGAAGTTTGGCAGGC |
| *KCNV1* | NP_055194.1 | 0.29 | 4.06 | *kcnv1* | XP_688404.1 | 0.6558 | 001 | 5' | GTCGGGACAAGTACTACCGCAGG | AAGCCGCTACATCCTCTCTC | CCATGTTCACGATGGACACC |
| | | | | | | | | 3' | ACATATCTGAGGCTGAGCTGCGG | TCTTTGGGCTTGACGATTGC | ACCAGAGATCTCCTGCACTG |
| | | | | | | | 201 | 5' | GTAGGTTTATAAGCTGAAGGTGG | AGCAGCCTTAAAATGTCTCCTG | ACTGAGAATTTCCAAGTGGTCC |
| | | | | | | | | 3' | AGCTGCACTGAGGCATGGGGAGG | | |
| *KDM1A* | NP_001009999.1 | 0.28 | 4.68 | *kdm1a* | NP_001229924.1 | 90.34 | 001 | 5' | ACCTGGTTCTCCACTTGGGGAGG | CTGGCCTCTGGTTGTCTATTG | CCACAATCGTGCCTGTTCTC |
| | | | | | | | | 3' | TCTGTTGACATGGCAGCCGAAGG | CTACACATTGTCACTGCGGC | GCATCCACTTCTACTGCGTC |
| | | | | | | | 003 | 5' | ACCTGGTTCTCCACTTGGGGAGG | | |
| | | | | | | | | 3' | CTAAATAGCTGTGTATCCGGTGG | ACAACCCCAAAGTCCAGCTC | ACCATAGCACCCAAGTCAGC |
| | | | | | | | 004 | 5' | GGTGATCGTGATTGGTGCCGGGG | | |
| | | | | | | | | 3' | GAAAGTTGAAGTCAAGCTGGTGG | GTATCACCAGGAAACAGCAAAG | GGCAAACAGAGTTTACTCACTG |
| *KIF1B* | NP_001352880.1 | 0.17 | 3.60 | *kif1b* | XP_009295087.1 | 80.87 | 201 | 5' | GCCATGTGCACCAAACACAGAGG | AGGCCATGCTCAAGGTAAAG | AATAACAGCAAAGTTGGCGC |
| | | | | | | | | 3' | GGTTTTGTGACGCAAAGCTGGGG | CAGCTGTAATTAATCACCCTCCC | CGCTGTGATGTTACATGCATAC |
| *KLHL11* | NP_060613.1 | 0.49 | 3.18 | *klhl11* | NP_956077.1 | 74.69 | 001 | 5' | CTCCGAGCAGTGAGTAGCGGAGG | CACTTCGTGAACATTGGCGG | GTTTGAGTGTGCTCCACCAC |
| | | | | | | | | 3' | TGTGTGGCAAATTGACCCAGCGG | TGCCACTTGTCTTGATCGGG | AGCCTGTCTACAGCTTGTGTC |
| *MAST1* | NP_055790.1 | 0.16 | 5.77 | *mast1a* | XP_009292706.1 | 78 | 001 | 5' | TCACCAATATGTCCTGGCAGAGG | CAACTGCATTTGTGTGTGTGTG | AGTGGGCTGCTTCCTATGAG |
| | | | | | | | | 3' | ATTACTGACAGTAAGAGGCGCGG | CGCAGGTCCAGTTACAAGTC | TGGGGTACTTGAAGCAGGAG |
| | | | | *mast1b* | XP_009297388.1 | 71.93 | 001 | 5' | TACTGGGAGAGAAGTTACGAGGG | ACGCAGGTTAGGGTAATTAGGC | GTGTGTGTGTGTGTGTGTGTG |
| | | | | | | | | 3' | CGTTGGGTGAAACAACCCAGAGG | ACCCTCTCCACCTGATCAAAC | TGACTTGCTTGCCTCTGGAG |
| | | | | | | | 201 | 5' | TGTCGTAGCAGGAGTGAGAGCGG | CCAACCCCTGTATGTGTTGC | GTAGAGGTGTGAAATCAGGTAGAC |
| | | | | | | | | 3' | CGTTGGGTGAAACAACCCAGAGG | | |
| *MED12* | NP_005111.2 | 0.07 | 5.51 | *med12* | XP_009289312.1 | 72.89 | 001 | 5' | ATTTCTGGCTCGTCACAGCGAGG | TGAGTTAACCGCATTGAACGTC | TTTTGCACCATGGACGAAAGC |
| | | | | | | | | 3' | CTCACCATGGGCTGTTGCGGGGG | GTTTGTCCACACAGCCACAG | ACTTTTTGGCCCATGAGGAGG |
| | | | | | | | 002 | 5' | TTCATCAATGCAGCCAGCACAGG | ACGTAGGCACCATCTAGCTAC | GATTGTGTGCCCATATGCCG |
| | | | | | | | | 3' | GAGTTGCCTGACCAGTGCTGCGG | | |
| *MYT1L* | NP_001289981.1 | 0.09 | 4.77 | *myt1la* | NP_001038364.1 | 68.09 | 001 | 5' | GCCTAATCGCACCACACAGGAGG | AGGCCCTGCTAGTACCTTTG | TTAACCCTCTCAAGGCAGGC |
| | | | | | | | | 3' | TCGGTTTCTATAGAAGGGAGAGG | CATTTCTTGCGGTGCACCTC | GGAGGACTAGGGGAGGAAAATG |

| | | | | | | | | | | | |
|---|---|---|---|---|---|---|---|---|---|---|---|
| *NBEA* | NP_056493.3 | 0.07 | 5.51 | *nbeaa* | NP_001013518.2 | 83.25 | 001 | 5' | ATCGAAGGCCGGGATCATGGAGG | CCCAGACGTTTTAAGGCCATC | AAAGGATTGATGCGCCTCTG |
| | | | | | | | | 3' | TGATGCCTGCATCACAGCCGGGG | GGATGCATTTGTTCCCTTCAG | AGCCTCAAGTATAAACCAGGC |
| | | | | | | | 201 | 5' | GGTTGCGGACGCTCTTGCGCAGG | CTCTTTCTGTCTTTGTCAGCTGG | CAGTAAAGACGTCTGTTTCCAGC |
| | | | | | | | | 3' | CTGGACACTGAGATCAGACGGGG | GCTGCTGTTTTTGTGTATTGAGCC | TGCAGTGTGTGTGTGTGTGTG |
| | | | | *nbeab* | XP_017206693.1 | 81.66 | 001 | 5' | AATGGCACAAAGACATGGACCGG | CCTGTGGGATTTGTGTCATTTGG | GTGCATTGTGGGTAATAATTGGGG |
| | | | | | | | | 3' | TGCATCATCTACTTTGACCGAGG | CCTCCACAAGCATGGCAATAC | GCACTGTGTGAATGGAGCAC |
| *NEDD4L* | NP_001138439.1 | 0.20 | 3.73 | *nedd4l* | XP_021324687.1 | 78.66 | 001 | 5' | TTTGTACTAAGGCAAGCTCGCGG | GCCTTGCCATTCTAAGGTCTG | TGTGCCTTCAGTAGATCCAGAC |
| | | | | | | | | 3' | CCAGACCACACATCAACAGCTGG | AGGGTGATCTAGAGGATGATGAG | GCTGGAAACCTGTAACCATTG |
| *NIPBL* | NP_597677.2 | 0.03 | 5.57 | *nipbla* | XP_009299463.1 | 74.03 | 001 | 5' | GGTGGTGCCCGGTAAAGGGGAGG | GCGGCTGTTCGATTTCTACC | AACAAGCATGCAGAGGTGTG |
| | | | | | | | | 3' | GCAGGAAAAGGAGACGACGGCGG | AGGACGCTAACAATTCAGGAGG | ATCATCGTCATACTCCTCATCTCC |
| | NP_597677.2 | | | *nipblb* | NP_001154919.2 | 59.42 | 201 | 5' | GGTGGTGGCAGGTAGAGGAGAGG | CGGATCGAAATGGGAAGTCTTTG | AAGGCCATAAGGTGAGGATGTC |
| | | | | | | | | 3' | GCGTTGGCGAAGTCCAGCAGGGG | AGTAGCAGCAGCAGTAGCAG | TCAAAGCTGATTGCACCAGAAC |
| *OGDH* | NP_002532.2 | 0.27 | 4.76 | *ogdha* | NP_957073.2 | 86.59 | 001 | 5' | TCTGTGAGGCCGTTATAGGCCGG | AAGACGGGAGCTGACACTTC | TCCAACTCCATGTCCTTCGC |
| | | | | | | | | 3' | CAAGAGGAACATAAGAACCAAGG | TTGTCTGACTGATGCTCTCGC | GTGTTCTTGTTTCCGGTGGC |
| | | | | | | | 003 | 5' | GCCAGGCTCACACCCAATGGAGG | AGTCCTTGCTCCCTTTCTGC | GGTTAAAGGCAGCACTGAGATATC |
| | | | | | | | | 3' | CCTGACTTACCTCTAAGCGACGG | TTTACCCTGTCCCCCTCTTC | GGTGAACCTAACCCATTAAGCTAC |
| | | | | *ogdhb* | XP_001338181.2 | 84.13 | 001 | 5' | CTCACAACACAGGCCGGCGGTGG | GGGTAAAGGAGTTCAAGCACTTTC | TTGTGCACGCTCTTGGGATC |
| | | | | | | | | 3' | GCATCTGCTCTGGTGTCAGGAGG | AAATCAGAACCACTGGCCCG | GGAGGGCGAATACTTTTGACTTC |
| *PACS1* | NP_060496.2 | 0.18 | 3.71 | *pacs1a* | NP_001092218.1 | 67.22 | 001 | 5' | GATAAACCGACCGACACCGGCGG | TCTCTAGCGGTTCACAAGCC | GCTGAAAACCCACGGTCTAAAG |
| | | | | | | | | 3' | CCGTCACCATCACTAAGCGGAGG | AATACTCGCTTTAGGTGCAGAC | TCTTCTCCTTGGTCACCACAG |
| *PC* | NP_001035806 | 0.06 | 3.76 | *pcxa* | NP_001315285.1 | 83,6 | 201 | 5' | TGAAGGTAAGCCGCTACAGGTGG | ACACACTGATCGAATGCACTG | TTGTAGCTCCCCATGTTTTACC |
| | | | | | | | | 3' | TGGGAGATCTTAATAAAGCGGGG | TGCTGAAGAGTTTGAGGTGGG | CACGAGCAAAGAGATTCACTGG |
| | | | | *pcxb* | NP_571625.2 | 83.62 | 201 | 5' | CCAGGGTTTGGGAAGACAGGTGG | ACCATGACACACTTGAACCAG | AGCACTAATCCAGGGATCAGTC |
| | | | | | | | | 3' | CCTGTGACAGAACCAGACCAAGG | GGTTTCCTCAAACGATTTGGG | AAACAATCCAATCTCGACGGG |
| *PHIP* | NP_060404.4 | 0.11 | 5.14 | *phip* | XP_009295279.1 | 66.27 | 001 | 5' | ACCGTGTCAGGACGCAGCAGAGG | CATCTGCCTTTCCCACAACG | GCACAAAAATCGCACATTACGG |
| | | | | | | | | 3' | CAAGCCCCAAAACTCTCCGGCGG | AAATAGAGACGCCGGTGGAC | CTTTGCTGGACGTCCCAAAAG |
| *PRPS1* | NP_002755.1 | 0.38 | 3.73 | *prps1a* | NP_001346823.1 | 96.25 | 001 | 5' | GGGGAAGCAAGGAATGACCGCGG | GTGCACTTATCATCTTGAGTGACG | CTTACAGAAAGGAGCCCACATG |
| | | | | | | | | 3' | CTGCAGCAAGATTCAGGCACAGG | CCTGCTAATCAGTCATGCTTTC | ATCTAGAATTCATGACCCAGCC |
| | | | | *prps1b* | NP_001070036.1 | 96.86 | 001 | 5' | GTCGGGCATAAGGAAAGCATGGG | GCATGCGTTTTATTTGTCGTGG | GGTTTGGCTATAGACATTACCAGC |
| | | | | | | | | 3' | GCCATCCGCAGAACCCACAACGG | CTATTGGCAGTTGAGACGAGC | AGGTTGACAAACGTTTGCCTG |
| *RALGAPA1* | NP_001333177.1 | 0.23 | 3.53 | *ralgapa1* | XP_021322984.1 | 68.84 | 002 | 5' | AGCACTGCGGATAAAGGCTGAGG | CTGCCCCATTTCCCTTTCTC | TGTGTCAGGATTAGGCTTGC |
| | | | | | | | | 3' | TTTTAGGGCATGACGAGGCGCGG | AGCTTTTCACGGTGGTCTTG | GCACATCACTTGACTTTTCCC |
| | | | | | | | 003 | 5' | TCGTGAAACACACAAAATCGCGG | TGGAAAGGTGTTGTAGGGGC | GCTTCAGGAGAGGTGGATAAAATG |
| | | | | | | | | 3' | CGCAGGACAAAAGACACGAGCGG | | |
| | | | | | | | 201 | 5' | TTGTAGGTCACAAATCCCAACGG | CCTGTGGGCTTTTTGAGATCC | ATGGTGGAAGGGAGTGACAC |
| | | | | | | | | 3' | CGCAGGACAAAAGACACGAGCGG | GCTGTTTTGAAATCGATGCACC | AATTGTGAATGGTGAGCAGTGC |
| *RBM8A* | NP_005096.1 | 0.56 | 2.16 | *rbm8a* | NP_001013363.1 | 93.55 | 001 | 5' | CCTATGGACGAGGATGGCGATGG | TCTCACTCACAAGCAAGCATC | GAGTATTGCACACATTCACCAC |
| | | | | | | | | 3' | GGCTGGGAAACAGACACCGGAGG | GGATTTGGTAATGATACAGGCCG | ACATGTACGTGTCAGCAACTTC |
| *RHOG* | NP_001656.2 | 0.89 | 2.49 | *rhoga* | NP_001076273.1 | 73.3 | 001 | 5' | TTCAGCGCTCAGCCAGGACGGGG | TGTTTGAACACTGCCCGAATG | GAGGACGGTTGTTTGGGATG |
| | | | | | | | | 3' | TTACAGCGCACAAGTGCTCGTGG | ACTTTGCGTATGTGTACCTTGC | TACGCAAACGGTCGTACTCC |
| | | | | *rhogb* | NP_956334.1 | 80.1 | 001 | 5' | TCTTGAGGACAGTAATCCGGCGG | TCGCGTCCTACAGACTTCAAC | TCCGAATTATTGACGCGCTTC |
| | | | | | | | | 3' | GAGGGTAGGACAGCGTGCGAAGG | TGCCTGCTCATCTCCTACAC | CTCTTGGTTCCCACAAGCAG |
| *RUVBL2* | NP_006657.1 | 0.11 | 3.11 | *ruvbl2* | NP_777285.1 | 88.34 | 001 | 5' | GGCAACCACAAAGGTTCCGGAGG | CCTAACGTCTATTGAGCGGTC | ACCATGTTGCCCCCATTTAG |
| | | | | | | | | 3' | GGGCACAGAAGTCCAAGTGGAGG | CCGTAAACGCAGGGTAATGTC | CTCTTTGCTGCAAGATCTCCAG |
| *SLC12A5* | NP_001128243.1 | 0.14 | 4.70 | *slc12a5a* | XP_021332993.1 | 77.74 | 001 | 5' | TGAGGAATAGAATCACACCCAGG | AAACCAATGCAATGTCCCCAC | TGAGACGCAGCCTTTTAATGAC |
| | | | | | | | | 3' | GGACCACCCCGAAACAGAAGCGG | GAACACTCTCTGACGGATGC | CTGAGGTGGTTTATGAGGATACAG |
| | | | | *slc12a5b* | NP_001289172.1 | 74.53 | 201 | 5' | TATGAGTCTGAAACTAACCGTGG | TTACACGGTGTCCTGATTACTG | GGATCTGAAGAAAAGTGTGGC |
| | | | | | | | | 3' | GAAGGCCAATAACCCAGCAGTGG | GTCAACCTTTTCCACCCTCTC | CTGCCAGAATAAGAAAAGCAGC |
| *SLC20A2* | NP_001244109.1 | 0.33 | 2.63 | *slc20a2* | XP_005167225.1 | 77.22 | 001 | 5' | TTCGGTGGGAGCCAATGACGTGG | ACGACCTTCATAGTACAGCACC | TTCCCTTCCGGATGGTTTCC |
| | | | | | | | | 3' | AATAGTGAATCCACTGCGGGAGG | CTGCATGCTGTGTGTGTTTACC | AAAACAGACGTCCCCTCTCG |
| *SLC2A1* | NP_006507.2 | 0.24 | 2.93 | *slc2a1a* | NP_001034897.1 | 74.39 | 001 | 5' | AGTGTGGTTAGAGTGGTCGGCGG | GGGATGCAGAATAAGATCATGC | AGTATCGGTAAGGTTCTGACTG |
| | | | | | | | | 3' | GGAGAAACCAGCGACAGCGAAGG | GGCACTACTGGTATGTGAACAC | AGATGTGCTCAAGTTTCCGTAC |
| | NP_006507.2 | | | *slc2a1b* | XP_002662574.1 | 79.67 | 001 | 5' | TGACAAGACGTGATGGAAGGCGG | TCCGAGGAAATCCAAGTGGG | CCACACCATGCTCTGTTACG |
| | | | | | | | | 3' | TGATGCCTGACGGAAACCGGCGG | CCTCTTTGTCCATGTGTTGAAC | GGCGAGTGCAGAAATGTTTG |
| *SMC1A* | NP_006297.2 | 0.06 | 6.45 | *smc1al* | NP_997975.2 | 89.78 | 001 | 5' | CCGCATCAATAATAAAGTGGTGG | ACAGAGCGTGGTGGTTTTTG | GCTAGCAACATCTCTTCCACTATC |
| | | | | | | | | 3' | AGTTTGAAGCTGCTAGGAAGAGG | CAGACAGTGTGGACAAGACATG | CTGTTAACCGTTGACTTCCATAGG |
| *SMC3* | NP_005436.1 | 0.04 | 6.40 | *smc3* | NP_999854.1 | 95.15 | 001 | 5' | GTCGAATATGATTTCCACGAAGG | TCATTGTCACTACCCATACGGAAC | TAAAGACGAGGTCACGATAAGGC |
| | | | | | | | | 3' | AGGCAGGACGAGCTGGACAGAGG | GAAAGCTGGAGCAGTGCAAC | TTGAAGGTGAGCTGGATGGC |
| *STT3A* | NP_689926.1 | 0.49 | 3.62 | *stt3a* | NP_958866.1 | 96.45 | 201 | 5' | GCTTGGTCATCTTTACACACGGG | AACCGCTGTTGTCTGTTTTGTC | AGAGGAATGCTGATGTTCGGG |
| | | | | | | | | 3' | GTCCACACGAAACTCTCCAGTGG | TTCATCTGATGGTACGGGATG | ATGAACTCACTGGCTTCGG |
| *STT3B* | NP_849193.1 | 0.25 | 3.76 | *stt3b* | NP_001315090.1 | 89.48 | 001 | 5' | GCTGGCTCTTTTGACAACGAGGG | TCTTCAGTGGCTTGACTGC | AGACAAACACAAAACGGTGC |
| | | | | | | | | 3' | AGACAGCCAAGAGGAAGCGGGGG | GCCCCTTGGAAAAATGAATTGC | TCCGTTCTCAACATTCGCATTC |
| *STX1B* | NP_443106.1 | 0.27 | 2.94 | *stx1b* | NP_571598.1 | 96.88 | 001 | 5' | TAAGGACAGTGACGATGACGAGG | CGGAAAACTTACTGCTGTTCAC | TCACAGCATAACCTGATGACC |
| | | | | | | | 001 | 3' | TGTGGACTATGTAGAGAGAGCGG | ACACGACGATTGTTTGATGTGC | TGTGTTCTCCAGCAAGGTGC |
| *STXBP1* | NP_003156.1 | 0.09 | 4.26 | *stxbp1a* | XP_005161170.1 | 88.06 | 001 | 5' | TAGTGGACCAGCTCAGCATGAGG | | |
| | | | | | | | | 3' | CTGCAGTCGACTCTGCGCAGAGG | | |
| | | | | | | | 002 | 5' | GACTGCAGAAGTGCTGAAGGAGG | | |
| | | | | | | | | 3' | CGAGTGAGTAGAACCTGCGGAGG | | |
| | | | | *stxbp1b* | NP_001082845.1 | 78.31 | 001 | 5' | AGACAGGTTACCTCATCAGATGG | | |
| | | | | | | | | 3' | TATGAGGTCACGCAGGCCAACGG | | |
| *TAF1* | XP_005262352.1 | 0.04 | 5.49 | *taf1* | XP_021331844.1 | 76.3 | 001 | 5' | TGTGTACCTGTTCTCCGCGAGGG | GTGTGCTAATTTTGTGTGTGGG | ATTGGTTGAGTAGGGGGAGG |

| | | | | | | | | | | | |
|---|---|---|---|---|---|---|---|---|---|---|---|
| [illegible] | [illegible] | [illegible] | [illegible] | [illegible] | [illegible] | [illegible] | [illegible] | 3' | ACGGATTGGACTTGAACAGGAGG | GGCGATAATCCATTCTCCTGTAAG | GCCAATGACATGACCACATTG |
| *TBL1XR1* | NP_078941.2 | 0.11 | 4.20 | *tbl1xr1a* | XP_005172880.1 | 96.11 | 001 | 5' | GACGCCAACAAATTACCAGTCGG | CTACAGTCTGCCAAAATGAGGG | TTAGACCCCCAGATCAGACG |
| | | | | | | | | 3' | CAACCAGCGCTAACATGCAGGGG | TTCTTTCTGCAGGACGGCAC | AAACCGCAGACTAGAGCTGAC |
| *TBX1* | NP_542378.1 | 0.43 | 0.74 | *tbx1* | XP_005168668.1 | 64.35 | 001 | 5' | TGATTGTCACAAAAGCCGGGAGG | AACGCTGGTGAAAAAGAACCC | GCATTGACGATGAAGATAGGCTAC |
| | | | | | | | | 3' | CTGGGTGAGGGATAGGGGGACGG | AATGTCTCGAGTCCTCAGCC | CATAATTAGTCGGTGCGCTCG |
| *TFAP2A* | NP_001358995.1 | 0.26 | 2.59 | *tfap2a* | XP_005162615.1 | 85.45 | 203 | 5' | TTGGGCGTATGAGACAGCGGAGG | GACACTTTATCGTTCTCTCTCTCC | GAGAGTAAGGGTCCTGAGACTG |
| | | | | | | | | 3' | TCTGCAGAACTATCTGACGGAGG | ACGTCCACAACCCATTCTCG | CTTTGTCACCCGCTTTCGAC |
| *THOC2* | XP_016885151.1 | 0.08 | 5.53 | *thoc2* | XP_005157191.1 | 70.26 | 201 | 5' | TTTTTAACCAGTCAAACGGGAGG | ACGGCTGCTTCATCTGAGAG | ACCCGTGTTCATTCCACCTG |
| | | | | | | | | 3' | CCAGGGACCGCTCCAAAGAGAGG | GCTAATGAAAGGCAGCAGGC | AGCCTTGTAAAATGTCCTGCTC |
| *TRIM8* | NP_112174.2 | 0.23 | 2.74 | *trim8a* | XP_009305420.1 | 66.43 | 201 | 5' | TGATCGGGTGAAAGACATCGAGG | AGGTCCCATTTCTGATCTGTC | AATGTCACGGCAAAAGAAGC |
| | | | | | | | | 3' | TGGATTGGAATAGTCGGCGGAGG | TCCCAGAAAGCAGCACTAGC | AGTGCGGTGGTTGGATATGG |
| | | | | *trim8b* | NP_001038379.1 | 66.55 | 001 | 5' | GCAAATGCTGATTAAACAGCAGG | ACTTTCGAAACAAGCGAAGGG | TGGCTTGTCAGAGTTTCACAC |
| | | | | | | | | 3' | ATGGGAGTGTGGGAACACCGAGG | TCCGCATCAGATTCAGCCTC | GCTGTTTCTGGTGAAGCTGC |
| | | | | | | | 002 | 5' | GCAAATGCTGATTAAACAGCAGG | | |
| | | | | | | | | 3' | AGGTGCTGCACTTTGTCCTGAGG | TAGCGGTGAGTTCGGTGAAC | GCTGTTTCTGGTGAAGCTGC |
| *TRIO* | NP_009049.2 | 0.14 | 5.32 | *trioa* | NP_001097996.1 | 82.36 | 201 | 5' | TCACGATGTAGTCCAGCAGACGG | | |
| | | | | | | | | 3' | CTGGTCAGAGGGAAGGGCAGGGG | | |
| | | | | | | | 202 | 5' | TCACGATGTAGTCCAGCAGACGG | AGGCAAGCCTAATGGAGACAG | GGAACATCCTGTGTAATGCTGAC |
| | | | | | | | | 3' | CTGGTCAGAGGGAAGGGCAGGGG | TGAAGAAGAGGGTGATAAAGGCTC | TTACCTTCTCAGCGTCCTGC |
| | | | | *triob* | XP_009290354.2 | 79.2 | 002 | 5' | TGCTGGATGGCCATAGGGGGCGG | GAACGAAGGATTGAGCAGTGG | ATGTCATTCGGCATCCACAG |
| | | | | | | | | 3' | GGTGGAAGAGATCGGCGTGGAGG | TAGGCCACGGTGTTCTGAAG | ACAAACAAGCGGTACCTGGG |
| *TUBB5* | NP_001280141.1 | 0.29 | 5.63 | *tubb5* | | | 001 | 5' | GTGAAGGCAAACCTGTGCAGAGG | GTCAAACTGGCGGACTTTGG | GTCTGACTGAGAAGGCGACTAC |
| | | | | | | | | 3' | CGACGCCAAGAACATGATGGCGG | TGGCGACCTCAACCATCTTG | ACATAGCCGTGAACTGCTCG |
| *U2AF1* | NP_001020374.1 | 0.22 | 3.85 | *u2af1* | NP_803432.1 | 95.19 | 001 | 5' | CCATTTTGAAAGAAGAAAGGCGG | TTGACCCTCCATCTTCAGCG | TAATGCTCAGGGGTGCTCAG |
| | | | | | | | | 3' | TGTACGGCCGCAGAAGGAAGAGG | GGTTCAGGTGAAGATGTGTGC | AAGGCATTATGACACCAGTGTG |
| *UBA1* | XP_016885266.1 | 0.11 | 3.48 | *uba1* | NP_998227.1 | 78.85 | 001 | 5' | ACGCGACGCTTCTTGGACAGCGG | AACTCGCTTTCAACTCGCATG | TTCAAGCCCCAGTGGTGTTC |
| | | | | | | | | 3' | AGTTGCTGAAGATCGTTCAGGGG | AAGCTGTCCTGTGAGCTACG | AGCTTACCTTGTGCTTGGGG |
| *UBR5* | NP_056986.2 | 0.07 | 6.21 | *ubr5* | XP_021322301.1 | 88.28 | 201 | 5' | GAACCAGACAGGAACCAGGGCGG | AGGGTGTTGCCGTGTTAGTC | CTGCATGAGTGGCTGTCAAC |
| | | | | | | | | 3' | CGCTCTGGAAGATCTGACAGCGG | ACGGGAAGCGTTATGAGCAC | CCCATGTCATTCATCTTGCGC |
| *USP7* | XP_016879141.1 | 0.06 | 5.65 | *usp7* | XP_009297739.2 | 93.78 | 001 | 5' | CAATGGGAACGTGGCTATGGCGG | GTGGAGCCATGTGTCTTACTTTC | CTTCAGAACGCCAGCTTGTG |
| | | | | | | | | 3' | ATCCGCATTTATTTTGGCAGGGG | GGACAGGTGCCATTGGTTTG | ACAGGTTCAGAACAAGTGGAGAG |
| *WDR82* | NP_079498.2 | 0.40 | 3.29 | *wdr82* | NP_001159740.1 | 93.61 | 001 | 5' | GCTCTAACAAGATTGACGGTAGG | AAATAATCGGCCGCCTCCTG | TGGAATTTGCTGGGGGAAAAAC |
| | | | | | | | | 3' | ACATTTCCAGGATCAGAAGATGG | AGGCTTCTTTCACACCAGACTC | AGGCACTGGCAAAGGTCATG |
| *WHSC1L1* | NP_075447.1 | 0.14 | 3.73 | *whsc1l1* | XP_021335001.1 | 62.05 | 001 | 5' | TGGTGGTGGTGAGGTAACAGAGG | CAGTTTGCTGCTTGGTGAGG | AATGGGGTTGGAGATGGTGC |
| | | | | | | | | 3' | GGTCCTGAAAAAGCACATGAGGG | TGGAGTACGTTGGCGAACTG | AGGTGAAGAGTCCAATCCGC |
| *WNT8B* | NP_003384.2 | 0.49 | 3.02 | *wnt8b* | NP_571034.1 | 81.95 | 001 | 5' | ACCCACAGCGGACTCAGAAGCGG | TCCTTTGGTGTGGTTTGCATTG | GGGCACATCAACATCGTGAG |
| | | | | | | | | 3' | AAAAGAGAAGTACCACCGGGCGG | TGCCTCCAGAACTGAGCATG | TAGAGTGCGGTTCTCCAAGC |
| *XPO1* | NP_003391.1 | 0.05 | 6.01 | *xpo1a* | XP_009292977.1 | 92.72 | 001 | 5' | TGTCTCCTTCAGCTCTATGCCGG | ACTCTACCGTTGCCTTGAGTG | TGGTCCTTCAGGTTGGTGAG |
| | | | | | | | | 3' | CGGTCTTGAGGAACTTCCAGTGG | ATGCAGCATCAGGGAACAGG | ATTAACTGCACAGCCCTCGC |
| | | | | *xpo1b* | XP_009291328.1 | 96.27 | 001 | 5' | TCAGCAAAGAATGGCCCAGGAGG | CTCAGCCAGTATATTGTTGCACC | AACCATGACCCACCAACTGG |
| | | | | | | | | 3' | GTACATGTTTAACCTCGTCGAGG | GGTCAGAATTTGCTTGGGTATG | AGCAAGTTGGCCACATACTC |
| | | | | | | | 003 | 5' | GGTGGAGAACGATCAGGGAGAGG | CCCATAAGTGAGTGTGTCGC | AGCATAGTCCAGATGGGTAAGG |
| | | | | | | | | 3' | TGCATGAGGAAGATGAGAAGCGG | GACTATGCTGACACAGAGCG | GGTAAAGTGTACCCTCTAGCAG |
| *XPR1* | NP_004727.2 | 0.16 | 3.23 | *xpr1a* | NP_001232029.1 | 79.83 | 001 | 5' | TGTTTCAGAGAAACTGGCCGAGG | ACAGTGGGAGCACAGTTTTG | GCCATGCTGCCCTTAAAAATC |
| | | | | | | | | 3' | GCAGAGGTCGACGTAAGGACAGG | CTGCAGATTTTCAAGTGGACTC | TGTGGCTCCTCTAAATGCTG |
| | | | | *xpr1b* | NP_001119862.1 | 79.08 | 001 | 5' | CCAAACGTGTGAGAAGGAGCTGG | ACTCATCTCTGGCTGCTAAC | TCAGCTTTGCTTCAAACACG |
| | | | | | | | | 3' | TAATTGTGGTGAGTTCAGAGCGG | AAATAGGTGGATGGAAACGCAG | GTCGTCTCAAGGACATGCTG |
| *YWHAG* | NP_036611.2 | 0.29 | 2.95 | *ywhag1* | NP_998187.1 | 95.95 | 001 | 5' | GACAGGGGAAAAGCGAGCCGCGG | GAAATGGTCCGCGCTTATCG | AGTAATTGAGCGCCAGACCC |
| | | | | | | | | 5' | GAGGGTGTCCAGCTCTGCGATGG | CATCTGGCGCTTAATTGTTGC | AGTCCTACAACGAGGCTCAC |
| *ZEB2* | NP_055610.1 | 0.11 | 3.94 | *zeb2b* | NP_001232895.1 | 70.33 | 201 | 5' | GTGTCCGGTGGCTTCAGGAGTGG | TTCATAAGAGAGGTATGCGCCG | TCGAGGATTTCTTCGGGAAGC |
| | | | | | | | | 3' | AGATGATATGGCCGACCCCGAGG | CCTGTGTGCTCATATTTGTGTCTG | TGAAGAGCTAAACGCCGCTG |
