## Supplemental Table 2 for "Zebrafish models of candidate human epilepsy-associated genes provide evidence of hyperexcitability"

| Gene | Mutation | Genotype | N | Paradigm | Larvae SII > 0 | Total SII |
|---|---|---|---|---|---|---|
| *abcc9* | NP_001025325.1: .pA1522RfsTer1551 | Ungenotyped | 192 | Light | 0 | 0 |
| | | | | PTZ | 0 | 0 |
| | | | | Spontaneous | 0 | 0 |
| *agpat3* | NP_998590.1: .pT35CfsTer58 | Ungenotyped | 284 | Light | 0 | 0 |
| | | | | PTZ | 0 | 0 |
| | | | | Spontaneous | 0 | 0 |
| *arfgef1* | XP_009295673.1: p.L367S_E368_E369del | -/- | 27 | Light | 1 | 6 |
| | | | | PTZ | 2 | 2 |
| | | | | Spontaneous | 2 | 3 |
| | | +/- | 41 | Light | 1 | 1 |
| | | | | PTZ | 0 | 0 |
| | | | | Spontaneous | 1 | 1 |
| | | +/+ | 35 | Light | 0 | 0 |
| | | | | PTZ | 0 | 0 |
| | | | | Spontaneous | 0 | 0 |
| *atp1a3a* *atp1a3b* | NP_571759.2: .pV943PfsTer950 | Ungenotyped | 192 | Light | 0 | 0 |
| | | | | PTZ | 0 | 0 |
| | | | | Spontaneous | 0 | 0 |
| *cacng2a* *cacng2b* | XP_056309537.1: .pQ305PfsTer326 | Ungenotyped | 192 | Light | 0 | 0 |
| | | | | PTZ | 0 | 0 |
| | | | | Spontaneous | 0 | 0 |
| *cacng3b* | XP_693299.1: .pF22Ter | Ungenotyped | 192 | Light | 0 | 0 |
| | | | | PTZ | 0 | 0 |
| | | | | Spontaneous | 0 | 0 |
| *calm2a* | NP_956290: .pA103GfsTer117 | Ungenotyped | 36 | Light | 0 | 0 |
| | | | | PTZ | 0 | 0 |
| | | | | Spontaneous | 0 | 0 |
| *chd2* | NP_001268391.1: .pF1752PfsTer1755 | Ungenotyped | 192 | Light | 0 | 0 |
| | | | | PTZ | 0 | 0 |
| | | | | Spontaneous | 0 | 0 |
| *cnnm2a* | NP_001138257.1: .pS36RfsTer144 | Cnnm2 -/-,+/+ | 59 | Light | 1 | 1 |
| | | | | PTZ | 0 | 0 |
| | | | | Spontaneous | 4 | 9 |
| | | Cnnm2 +/-,+/- | | Light | 0 | 0 |
| | | | | PTZ | 0 | 0 |
| | | | | Spontaneous | 0 | 0 |
| *cnnm2b* | XP_001922805.4: .pQ4PfsTer64 | Cnnm2 +/+,+/+ | 36 | Light | 1 | 1 |
| | | | | PTZ | 1 | 1 |
| | | | | Spontaneous | 2 | 6 |
| *csnk1g2a* | XP_005171263.1: .pH451RfsTer434 | Csnk1g -/-,-/- | 4 | Light | 0 | 0 |
| | | | | PTZ | 0 | 0 |
| | | | | Spontaneous | 0 | 0 |
| | | Csnk1g +/-,+/- | 10 | Light | 0 | 0 |
| | | | | PTZ | 0 | 0 |
| | | | | Spontaneous | 0 | 0 |
| *csnk1g2b* | NP_001039315.1: .pG32SfsTer125 | Csnk1g +/+,+/+ | 5 | Light | 0 | 0 |
| | | | | PTZ | 0 | 0 |
| | | | | Spontaneous | 0 | 0 |
| *cul2* | XP_002666627.1: .pD673EfsTer681 | Ungenotyped | 192 | Light | 2 | 2 |
| | | | | PTZ | 1 | 1 |
| | | | | Spontaneous | 2 | 7 |
| *dip2ca* | XP_009295792.1: .pA69VfsTer93 | | | Light | 0 | 0 |
| | | Ungenotyped | 192 | PTZ | 0 | 0 |

| | | | | | | |
|---|---|---|---|---|---|---|
| *dip2cb* | XP_698501.6: .pT1337GfsTer1451 | Ungenotyped | 192 | Spontaneous | 0 | 0 |
| *eftud2* | NP_956802.2: .pP76NfsTer112 | Ungenotyped | 192 | Light | 0 | 0 |
| | | | | PTZ | 0 | 0 |
| | | | | Spontaneous | 0 | 0 |
| *fgd1* | XP_688794.4: .pM163EfsTer181 | Ungenotyped | 96 | Light | 0 | 0 |
| | | | | PTZ | 0 | 0 |
| | | | | Spontaneous | 0 | 0 |
| *flnca* | XP_688794.4: .pM163EfsTer181 | Ungenotyped | 192 | Light | 0 | 0 |
| | | | | PTZ | 0 | 0 |
| *flncb* | XP_021330906.1: .pL2652TfsTer2557 | | | Spontaneous | 0 | 0 |
| *glud1a* | NP_997741.1: .pL18Ter | Ungenotyped | 192 | Light | 0 | 0 |
| | | | | PTZ | 0 | 0 |
| | | | | Spontaneous | 0 | 0 |
| *hcfc1a* | NP_001038529.1: .pR32TfsTer37 | Ungenotyped | 192 | Light | 0 | 0 |
| | | | | PTZ | 0 | 0 |
| *hcfc1b* | | | | Spontaneous | 0 | 0 |
| *hecw2a* | XP_003197660.3: .pR1308QfsTer1335 | Ungenotyped | 205 | Light | 0 | |
| | | | | PTZ | 0 | 0 |
| *hecw2b* | XP_003197660.3: .pR1308QfsTer1335 | | | Spontaneous | 0 | 0 |
| *hnrnph1* | XP_005169636.1: .pN377EfsTer404 | Ungenotyped | 96 | Light | 0 | 0 |
| | | | | PTZ | 0 | 0 |
| | | | | Spontaneous | 0 | 0 |
| *huwe1* | NP_001352583.1: .pfsTer4320 | Ungenotyped | 192 | Light | 0 | 0 |
| | | | | PTZ | 0 | 0 |
| | | | | Spontaneous | 0 | 0 |
| *iqsec1a* | XP_009304345.1: .pP355_R416del | Ungenotyped | 288 | Light | 0 | 0 |
| | | | | PTZ | 0 | 0 |
| *iqsec1b* | | | | Spontaneous | 0 | 0 |
| *kcnd2* | p.M375del | Kcnd2 -/- | 28 | Light | 0 | 0 |
| | | | | PTZ | 1 | 1 |
| | | | | Spontaneous | 1 | 1 |
| | | Kcnd2 +/- | 47 | Light | 1 | 1 |
| | | | | PTZ | 3 | 3 |
| | | | | Spontaneous | 4 | 4 |
| | | Kcnd2 +/+ | 21 | Light | 0 | 0 |
| | | | | PTZ | 0 | 0 |
| | | | | Spontaneous | 1 | 1 |
| *kcnj2a* | NP_001318145.1: .pV414PfsTer415 | Ungenotyped | 192 | Light | 0 | 0 |
| | | | | PTZ | 0 | 0 |
| *kcnj2b* | | | | Spontaneous | 0 | 0 |
| *kcnj3a* | XP_002663356.2: .pD64EfsTer98 | Ungenotyped | 192 | Light | 0 | 0 |
| | | | | PTZ | 0 | 0 |
| *kcnj3b* | XP_005165913.1: .pW315VfsTer362 | | | Spontaneous | 0 | 0 |
| *kcnv1* | XP_688404.1: p.H433Ter | Kcnv1 -/- | 27 | Light | 1 | 1 |
| | | | | PTZ | 0 | 0 |
| | | | | Spontaneous | 3 | 5 |
| | | Kcnv1 +/- | 106 | Light | 2 | 2 |
| | | | | PTZ | 0 | 0 |
| | | | | Spontaneous | 6 | 11 |
| | | Kcnv1 +/+ | 27 | Light | 1 | 0 |
| | | | | PTZ | 0 | 0 |
| | | | | Spontaneous | 0 | 0 |
| *kif1b* | XP_009295103.1: .pP64LfsTer73 | Ungenotyped | 192 | Light | 0 | 0 |
| | | | | PTZ | 0 | 0 |

| | | | | | | |
|---|---|---|---|---|---|---|
| | [illegible] | | | Spontaneous | 0 | 0 |
| *klhl11* | NP_956077.1: .pV462TER | Ungenotyped | 192 | Light | 0 | 0 |
| | | | | PTZ | 0 | 0 |
| | | | | Spontaneous | 0 | 0 |
| *mast1a* | XP_009297388.1: .pT1378GfsTer1379 | Ungenotyped | 192 | Light | 0 | 0 |
| | | | | PTZ | 0 | 0 |
| *mast1b* | XP_009292706.1: p.R1094_G195insVIEKRV | | | Spontaneous | 0 | 0 |
| *nipbla* | NP_001154919.2: .pfsTer85 | Ungenotyped | 96 | Light | 0 | 0 |
| | | | | PTZ | 0 | 0 |
| *nipblb* | | | | Spontaneous | 0 | 0 |
| *pacs1a* | NP_001092218.1: .pT394PfsTer403 | Ungenotyped | 192 | Light | 0 | 0 |
| | | | | PTZ | 0 | 0 |
| | | | | Spontaneous | 0 | 0 |
| *prps1a* | NP_001092218.1: .pT394PfsTer403 | Prps1 -/-,-/- | 6 | Light | 0 | 0 |
| | | | | PTZ | 0 | 0 |
| | | | | Spontaneous | 0 | 0 |
| | | Prps1 +/+,-/- | 91 | Light | 0 | 0 |
| | | | | PTZ | 2 | 2 |
| | | | | Spontaneous | 2 | 2 |
| *prps1b* | NP_001070036.1: .pfsTer95 | Prps1 +/+,+/+ | 50 | Light | 0 | 0 |
| | | | | PTZ | 0 | 0 |
| | | | | Spontaneous | 0 | 0 |
| *ralgapa1* | XP_021322985.1_: .pP2438AfsTer2445 | Ungenotyped | 192 | PTZ | 0 | 0 |
| | | | | Light | 0 | 0 |
| | | | | Spontaneous | 0 | 0 |
| *rbm8a* | NP_001013363.1: .pD22EfsTer25 | Rbm8a -/- | 51 | Light | 0 | 0 |
| | | | | PTZ | 0 | 0 |
| | | | | Spontaneous | 1 | 1 |
| | | Rbm8a +/- | 97 | Light | 1 | 1 |
| | | | | PTZ | 0 | 0 |
| | | | | Spontaneous | 3 | 3 |
| | | Rbm8a +/+ | 44 | Light | 0 | 0 |
| | | | | PTZ | 0 | 0 |
| | | | | Spontaneous | 1 | 5 |
| *slc12a5a* | XP_021332993.1: .pG124Ter | Ungenotyped | 190 | Light | 0 | 0 |
| | | | | PTZ | 0 | 0 |
| *slc12a5b* | | | | Spontaneous | 0 | 0 |
| *slc2a1a* | NP_001034897.1: .pP58LfsTer59 | Ungenotyped | 192 | Light | 0 | 2 |
| | | | | PTZ | 0 | 0 |
| *slc2a1b* | XP_002662574.1: .pE2LfsTer479 | | | Spontaneous | 0 | 0 |
| *stt3b* | NP_001038379.1: .pN203K_E204del | Ungenotyped | 192 | Light | 0 | 0 |
| | | | | PTZ | 0 | 0 |
| | | | | Spontaneous | 0 | 0 |
| *taf1* | NP_001038250.1: .pQ1796SfsTer1802 | Ungenotyped | 192 | Light | 0 | 0 |
| | | | | PTZ | 0 | 0 |
| | | | | Spontaneous | 0 | 0 |
| *tfap2a* | NP_001306087.1: .pS420IfsTer422 | Ungenotyped | 39 | Light | 0 | 0 |
| | | | | PTZ | 0 | 0 |
| | | | | Spontaneous | 0 | 0 |
| *thoc2* | XP_005157191.1: .pS488HfsTer533 | Ungenotyped | 192 | Light | 0 | 0 |
| | | | | PTZ | 0 | 0 |
| | | | | Spontaneous | 0 | 0 |
| *trim8a* | NP_001265783.1: p.P465del | Trim8 -/-,-/- | | Light | 0 | 0 |
| | | | | PTZ | 0 | 0 |
| | | | | Spontaneous | 0 | 0 |
| | | | | Light | 0 | 0 |

| | | | | | | |
|---|---|---|---|---|---|---|
| | | Trim8 +/-,+/- | 64 | PTZ | 0 | 0 |
| | | | | Spontaneous | 5 | 8 |
| *trim8b* | NP_001135847.1: .pH454IfsTer473 | | | Light | 1 | 1 |
| | | Trim8 +/+,+/+ | 90 | PTZ | 0 | 0 |
| | | | | Spontaneous | 2 | 2 |
| | | | | Light | 0 | 0 |
| | | Trio -/-,-/- | 1 | PTZ | 0 | 0 |
| *trioa* | XP_009292696.1: .pQ299AfsTer358 | | | Spontaneous | 0 | 0 |
| | | | | Light | 0 | 0 |
| | | Trio +/-,+/- | 12 | PTZ | 0 | 0 |
| | | | | Spontaneous | 2 | 4 |
| *triob* | XP_009290354.2: .pD1760EfsTer1869 | | | Light | 0 | 0 |
| | | Trio +/+,+/+ | 1 | PTZ | 0 | 0 |
| | | | | Spontaneous | 0 | 0 |
| | | | | Light | 0 | 0 |
| *u2af1* | NP_803432.1: .pR187QfsTer196 | Ungenotyped | 192 | PTZ | 0 | 0 |
| | | | | Spontaneous | 0 | 0 |
| | | | | Light | 0 | 0 |
| *uba1* | NP_998227.1: .pQ914PfsTer951 | Ungenotyped | 96 | PTZ | 0 | 0 |
| | | | | Spontaneous | 0 | 0 |
| | | | | Light | 0 | 0 |
| | | Ubr5 -/- | 14 | PTZ | 0 | 0 |
| | | | | Spontaneous | 0 | 0 |
| | | | | Light | 0 | 0 |
| *ubr5* | NP_001157866.1: .pL114RfsTer149 | Ubr5 +/- | 58 | PTZ | 0 | 0 |
| | | | | Spontaneous | 6 | 6 |
| | | | | Light | 0 | 0 |
| | | Ubr5 +/+ | 24 | PTZ | 0 | 0 |
| | | | | Spontaneous | 3 | 5 |
| | | | | Light | 0 | 0 |
| | | Wnt8b -/- | 44 | PTZ | 0 | 0 |
| | | | | Spontaneous | 0 | 0 |
| | | | | Light | 4 | 4 |
| *wnt8b* | p.S79_R80del | Wnt8b +/- | 93 | PTZ | 0 | 0 |
| | | | | Spontaneous | 2 | 2 |
| | | | | Light | 2 | 2 |
| | | Wnt8b +/+ | 51 | PTZ | 1 | 1 |
| | | | | Spontaneous | 1 | 1 |
| *xpr1a* | NP_001232029.1: p.E79_A80del | | | Light | 0 | 0 |
| | | Ungenotyped | 192 | PTZ | 0 | 0 |
| *xpr1b* | NP_001119862: .pF622RfsTer631 | | | Spontaneous | 0 | 0 |
| | | | | Light | 0 | 0 |
| *ywhag2* | XP_021328588.1: .pA206WfsTer228 | Ungenotyped | 120 | PTZ | 0 | 0 |
| | | | | Spontaneous | 0 | 0 |
| | | | | Light | 0 | 0 |
| *zeb2b* | XP_021332576.1: .pF245GfsTer302 | Ungenotyped | 176 | PTZ | 0 | 0 |
| | | | | Spontaneous | 0 | 0 |
