## Supplemental Table 3 for "Zebrafish models of candidate human epilepsy-associated genes provide evidence of hyperexcitability"

| Allele | Genotype | Behavior Paradigm | | | | | | | Local Field Potential | | | | Inhibitory Interneurons |
|---|---|---|---|---|---|---|---|---|---|---|---|---|---|
| | | Spontaneous | | | PTZ | | Light | | | | | | |
| | | N | SI | SII | SI | SII | SI | SII | Spike Rate | Spike Amplitude | Burst Rate | Burst Duration | |
| *arfgef1* | HET | 27 | ns | P(X=2)=0.13 | ns | ***P(X=2)=0.02** | ns | P(X=1)=0.23 | ns | ns | ns | ns | ***0.04** |
| | HOM | 41 | | P(X=1)=0.37 | | P(X=0)=0.69 | | P(X=1)=0.29 | ns | ns | ns | ns | ****0.009** |
| *kcnd2* | HET | 28 | | P(X=1)=0.36 | ns | P(X=1)=0.2 | ns | P(X=0)=0.73 | ***0.01** | ****.005** | ****0.009** | ***0.02** | ns |
| | HOM | 47 | ns | P(X=3)=0.1 | | ****P(X=4)=.001** | | P(X=0)=0.58 | 0.94 | 0.19 | 0.79 | 0.79 | ns |
| *kcnv1* | HET | 27 | ns | ***P(X=3)=0.03** | ns | P(X=0)=0.78 | ns | P(X=1)=0.23 | ns | ns | ns | ns | ns |
| | HOM | 106 | | ***P(X=6)=0.04** | | P(X=0)=0.38 | | P(X=2)=0.22 | ns | ns | ns | ns | ns |
| *ubr5* | HET | 14 | ns | P(X=0)=0.68 | ns | P(X=0)=0.88 | ns | P(X=0)=0.85 | ns | ns | ns | ns | N/A |
| | HOM | 58 | | ****P(X=6)=003** | | P(X=0)=0.59 | | P(X=0)=0.52 | ns | ns | ns | ns | |
| *wnt8b* | HET | 44 | 0.065 | P(X=0)=0.3 | ns | P(X=0)=0.67 | ns | P(X=0)=0.6 | ns | ns | ****0.008** | ***0.01** | ***0.01** |
| | HOM | 93 | ****0.0067** | P(X=2)=0.26 | | P(X=0)=0.43 | | ***P(X=4)=0.02** | ns | ns | ns | ns | ***0.03** |
